# A mechanistic framework linking within-host pathogen progression to vector-mediated transmission under climate forcing

**DOI:** 10.64898/2026.07.01.735761

**Authors:** Juan Carlos Rodríguez-Cabanillas, Manuel A. Matías, Àlex Giménez-Romero

**Author notes:** Corresponding author: Àlex Giménez-Romero.

## Abstract

Climate-driven disease forecasts typically assess whether environmental conditions favor pathogen growth, yet epidemic spread depends critically on how physiological processes within infected hosts shape transmission over time. This distinction is particularly consequential for vector-borne plant diseases, where vectors acquire infection from hosts whose pathogen load, symptom severity, and recovery are themselves temperature-dependent. Here, we develop a mechanistic epidemic framework that couples temperature-driven within-host pathogen dynamics to vector-mediated transmission. Infected hosts progress through ordered infection stages with stage-specific infectiousness, while transitions among stages—both progression and regression—are governed by thermal effects on pathogen accumulation and decay. We parameterize the model using experimental data for Pierce’s disease of grapevine, caused by *Xylella fastidiosa*, and analyze epidemic invasion under constant, seasonal, stochastic, and empirical temperature regimes. We show that temperature affects invasion not only by altering pathogen growth rates but also by reshaping the time hosts spend in transmissible infection stages. This generates a slow-growth paradox: temperatures that maximize within-host pathogen growth need not maximize epidemic spread, because rapid progression shortens the effective transmission window, whereas mildly suboptimal temperatures can prolong infectiousness and sustain larger epidemics. Conversely, cold conditions can suppress invasion by either halting progression or inducing regression and recovery. Analytical expressions for the basic reproduction number under constant and seasonal forcing capture these mechanisms and predict final epidemic size across diverse climatic regimes. Short-term temperature variability has its strongest effects near thermal thresholds, and empirical temperature series from invaded regions generate markedly different epidemic trajectories despite similar invasion suitability. These results show that ignoring the coupling between within-host physiology and transmission can qualitatively mislead predictions of plant disease dynamics under climate change, misidentifying the thermal regimes that pose the greatest epidemic risk.

## 1 Introduction

Mathematical models have long been central to the study of infectious disease dynamics. Since the foundational formulations of Kermack and McKendrick [1–3] and the subsequent development of epidemic theory in the twentieth century [4, 5], such models have provided powerful tools for identifying thresholds for disease emergence, elucidating mechanisms of epidemic spread, and designing effective control strategies [6, 7]. In plant pathology, modeling has become integral to risk assessment, surveillance, and management [8], particularly in agricultural systems where epidemics can have profound ecological and economic impacts [9, 10].

A pervasive simplification in most plant disease models is that hosts become infectious immediately after infection, or following a fixed latent period, and then transmit with constant infectiousness. Although mathematically convenient, these assumptions are often biologically unrealistic: they neglect the dependence of infectiousness on the age of infection—the time elapsed since a host was exposed [11]—a limitation already addressed by Kermack and McKendrick themselves [1]. In reality, infection unfolds through a sequence of processes, including pathogen establishment, tissue colonization, systemic spread, and symptom expression [12]. These processes dynamically determine pathogen load and, with it, the host’s capacity to transmit the disease. Infectiousness is therefore not a fixed property but an emergent, time-varying quantity shaped by pathogen-host biology. How long a host spends in each infection stage, and therefore how much transmission occurs before progression moves it onward, is as epidemiologically consequential as the transmission rate itself.

Environmental conditions compound this complexity by modulating the very processes that generate temporal variation in infectiousness [13]. Pathogen traits such as sporulation rate respond to temperature and humidity [14], with measurable consequences for disease spread [15] and establishment [16]. In vector-borne systems, insect population dynamics exhibit nonlinear thermal responses [17, 18] that alter transmission intensity, epidemic trajectories, and invasion thresholds [19, 20]. Climate thus shapes infectiousness through multiple interacting pathways simultaneously. Because temperature affects both pathogen growth and the progression of hosts toward advanced disease stages, the net effect of warming on epidemic spread is not a priori obvious and may be counterintuitive, a possibility that models assuming fixed transmission rates are structurally unable to capture [21].

Capturing this environmentally driven temporal variability remains a central challenge in plant disease modeling [22]. The dominant response has been to embed time-variation directly into model coefficients—seasonally forced transmission rates being the canonical example [23–25]. While flexible, this approach faces two interrelated limitations. First, fitting time-dependent rates requires extensive field data across environmental conditions and geographical contexts. Second, and more fundamentally, transmission rates are rarely measurable directly in natural systems [26, 27], so phenomenological time variation offers little mechanistic insight and limited transferability. The principled alternative—mechanistic models that derive transmission from experimentally measurable physiological processes—has proven difficult to implement, because translating controlled laboratory observations to the heterogeneous, fluctuating conditions of real ecosystems is far from trivial [22]. This difficulty reflects a deeper, persistent challenge in infectious disease modeling: despite broad recognition of the importance of coupling within-host dynamics to between-host transmission, quantitatively explicit frameworks that achieve this integration remain scarce [28, 29]. Consequently, the lack of a robust bridge between physiological measurements and field-scale epidemiology has limited the adoption of mechanistic, climate-sensitive models—precisely the class of frameworks increasingly called for as disease distributions shift under climate change [30, 31]. Without such a framework, predictions of how warming affects epidemic spread risk conflating faster pathogen growth with greater transmission.

Recent work on Pierce’s disease (PD) of grapevine has begun to close this gap. A climate-driven model was developed to parameterize population-level epidemic risk from experimentally measured pathogen growth rates and symptom progression data [32]. By explicitly linking temperature-dependent within-host processes—symptom development and host recovery—to between-host transmission, this framework provides a mechanistic route for translating controlled physiological experiments into epidemiological risk estimates. The framework successfully explained the current geographical distribution of the disease [32, 33], projected climate change–driven shifts in risk [34], reconstructed its biogeographic history [35], and identified Apulia, Italy, as a high-risk area before outbreaks were officially confirmed there [36]. These results have established a proof of concept: experimentally grounded, mechanistic linkage between within-host physiology and epidemiological risk is both feasible and predictive. However, those models were designed to assess only the suitability of disease establishment and relied on simplified epidemiological structures that cannot resolve detailed dynamics, such as transient epidemic trajectories, seasonal incidence patterns, or long-term prevalence.

Here, we develop a mathematically explicit framework that directly couples temperature-dependent within-host pathogen progression to vector-mediated transmission dynamics, resolving epidemic behavior well beyond the establishment threshold. The model partitions the infected host population into discrete infection stages, with temperature-dependent progression and regression rates that characterize the pathogen’s thermal niche [21, 37]. By linking pathogen load to host infectiousness at each stage, the framework translates controlled physiological measurements into time-varying infectiousness under fluctuating field conditions, directly addressing the challenge of environmental non-stationarity [22]. Applied to *Xylella fastidiosa* in vineyards (Pierce’s disease), the model yields a counterintuitive and analytically tractable result: faster pathogen growth does not necessarily increase epidemic spread. We show that this “slow-growth paradox” arises because accelerated within-host progression moves hosts rapidly to the chronic infected state, thereby shortening the effective transmission window and reducing the effective *R*_0_. These results indicate that conflating faster pathogen growth with greater transmission risk, a natural but unexamined assumption in models that lack within-host resolution, may lead to wrong predictions about how disease dynamics respond to climate warming.

## Methods

### The model

We extend a vector-borne epidemiological framework for host-vector-pathogen interactions [20, 32] to explicitly incorporate temperature-dependent pathogen progression within the host plant. This framework has been used to characterize *Xylella fastidiosa* diseases across various hosts and regions by parametrizing seasonal vector dynamics mimicking experimental observations [20].

To do so, the infected host compartment, *I*_*H*_, is structured into *n* ordered stages, representing increasing pathogen load and, consequently, increasing host infectiousness [21, 37, 38]. This construction is equivalent to the so-called Linear Chain Trick [39, 40], which replaces the memoryless (exponentially distributed) exit times of standard epidemiological models with a more realistic staged progression. Susceptible hosts, *S*_*H*_, become infected through contact with infectious vectors, *I*_*v*_, at rate *βS*_*H*_*I*_*v*_*/N*_*H*_. Once infected, hosts progress through infection stages at a temperature-dependent progression rate *γ*(*T* (*t*)) and may regress through them at a temperature-dependent regression rate *χ*(*T* (*t*)). Susceptible vectors, *S*_*v*_, acquire infection while feeding on infected hosts, with a stage-specific acquisition rate *α*_*i*_*I*_*H,i*_*S*_*v*_*/N*_*H*_ that reflects differences in within-host pathogen load. Note that the final infectious stage *I*_*H,n*_ differs structurally from intermediate stages: hosts there do not progress further, but are instead removed at rate Γ, and regression from this stage is not permitted, reflecting the irreversible nature of chronic infection. Infected vectors are assumed to remain infectious for the remainder of their lifespan; vectors are recruited and die at rates *δ* and *µ*, respectively.

This construction allows infectiousness to emerge from within-host pathogen progression rather than being imposed as a fixed trait. The full system is:

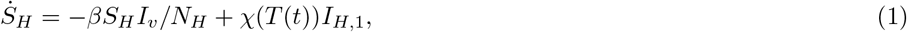

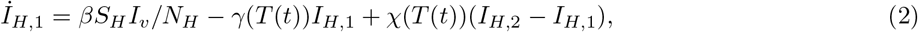

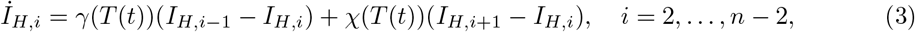

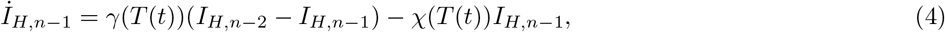

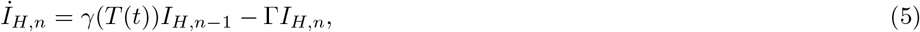

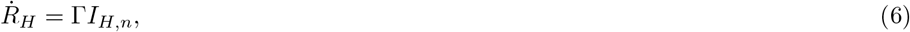

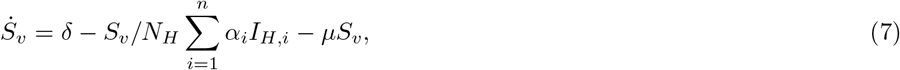

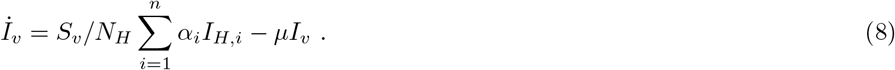

Table 1 summarizes the model variables and parameters. Note that the total number of infected hosts 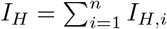 is the sum over all infected compartments, including the chronic compartment *I*_*H,n*_.

**Table 1.** Model variables and parameters.

| Type | Symbol | Definition |
| --- | --- | --- |
| Variable | $S_H$ | Susceptible hosts |
| Variable | $I_{H,i}$ | Infected hosts in compartment $i = 1, \dots, n-1$ |
| Variable | $I_{H,n}$ | Chronically infected hosts |
| Variable | $I_H$ | Total infected hosts |
| Variable | $R_H$ | Removed hosts |
| Variable | $S_v$ | Susceptible vectors |
| Variable | $I_v$ | Infected vectors |
| Parameter | $\beta$ | Vector-to-host transmission rate |
| Parameter | $\alpha_i$ | Host-to-vector transmission rate for hosts in state $i$ |
| Parameter | $\gamma(T(t))$ | Forward transition rate from infected-host compartment $i$ to $i+1$ |
| Parameter | $\chi(T(t))$ | Backward transition rate from infected-host compartment $i$ to $i-1$ |
| Parameter | $\Gamma$ | Mortality rate of chronically infected hosts |
| Parameter | $\delta$ | Vector birth rate |
| Parameter | $\mu$ | Vector mortality rate |
| Parameter | $N_H$ | Total host population |
| Parameter | $N_v$ | Total vector population |

### Modeling pathogen load dynamics

Local temperature mediates the growth and survival processes of the within-host pathogen population. We model the pathogen load, *P*, through a logistic growth law whose net rate depends on temperature-dependent growth and death rates, *f* (*T*) and *g*(*T*):

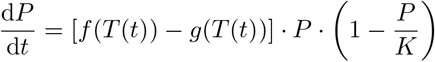

whose solution is

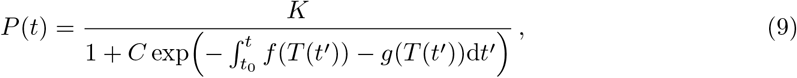

where the integration constant *C* = (*K* − *P*_0_)*/P*_0_ is determined by the initial pathogen load *P*_0_ = *P* (*t*_0_).

We define two thermal integrals that track the cumulative thermal drivers of pathogen growth and decay:

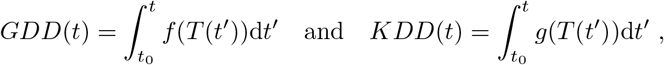

where *GDD* (Growing Degree Days) and *KDD* (Killing Degree Days) capture pathogen proliferation and die-off, respectively. Eq. (9) then becomes

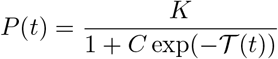

Where

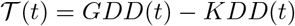

is the thermal integral obtained from the balance between pathogen growth (i.e., *GDD*) and decline (i.e., *KDD*). Thus, the thermal integral (*t*) shapes the within-host bacterial population and makes explicit that the pathogen load increases with accumulated *GDD* and decreases with accumulated *KDD*.

This formulation is general and does not depend on the specific thermal response of any particular pathogen. It provides a mechanistic link between environmental forcing and within-host pathogen load through the cumulative balance between growth and decay. Its application to a given disease therefore requires only the specification and parameterization of the functions *f* (*T*) and *g*(*T*), which encode the pathogen-specific effects of temperature on proliferation and mortality, respectively. These functions can be estimated from experimental or observational data and will generally differ among pathogens, host species, and environmental contexts [41–43].

### Temperature-dependent transition rates for Pierce’s disease

We followed the thermal-time parameterization previously developed for Pierce’s disease of grapevine [32], which we fully describe in Supplementary Section 1. The pathogen growth function, *f* (*T*), was derived from experimental measurements of the temperature dependence of *Xylella fastidiosa* growth [44], whereas pathogen decline was represented by the cold-response function *g*(*T*) = max(*T*_*r*_ − *T*, 0), with *T*_*r*_ = 6 ^*°*^C, based on empirical evidence for winter curing [41, 45] (see [46] for a review).

To link this continuous within-host process to the stage-structured epidemic model, we discretized the interval from initial infection to chronic infection into *n* = 150 infection stages. With *F*(*Τ*) denoting the empirical relationship between accumulated thermal time and the probability of chronic infection [32], we define *Τ*^*^ as the thermal accumulation for which

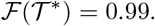

Thus, *Τ* ^*^ represents the thermal accumulation required for near-certain transition to chronic infection. The interval [0, *Τ* ^*^] was divided into *n* = 150 equal sub-intervals of width

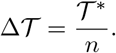

The temperature-dependent rates of progression and regression between adjacent infection stages were obtained by converting instantaneous thermal accumulation into stage transitions. Therefore,

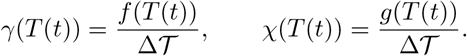

Here, *γ*(*T* (*t*)) moves infected hosts toward advanced infectious stages, whereas *χ*(*Τ* (*t*)) moves them backward. Because the experimentally derived function *F*(*Τ*) is a monotonic function of thermal accumulation, pathogen load increase with stage number. Therefore, stage-specific host-to-vector acquisition rates were assumed to also increase with within-host pathogen load [32]. For a host in stage *i*, with representative thermal load *Τ*_*i*_ = *i*ΔT, we set

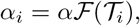

where *α* is the acquisition rate from chronically infected hosts.

### Computing the basic reproduction number

The basic reproduction number *R*_0_ is the expected number of secondary infections generated by a single infected individual in a fully susceptible population. Because *γ*(*T* (*t*)) and *χ*(*T* (*t*)) are time-dependent, the system is non-autonomous, and the exact invasion threshold requires a next-generation operator (in functional space) [47], a Floquet analysis in the periodic case [48–50], which does not, in general, yield a closed-form expression.

To obtain an analytically tractable approximation, we replace *γ*(*T* (*t*)) and *χ*(*T* (*t*)) by their averages over one annual cycle,

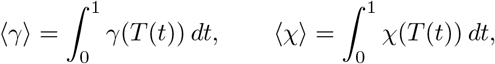

with which one can obtain the averaged Jacobian. This averaging is justified when the epidemic develops slowly relative to the annual cycle, so that infected hosts experience many seasonal periods, and the relevant quantities are the cycle-averaged rates 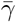 and 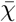 ; formally, it corresponds to the first-order term of the Magnus expansion [51].

Applying the next-generation matrix method [52] to the resulting autonomous system yields (see Supplementary Section 2 for the full derivation),

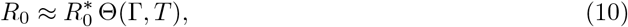

Where

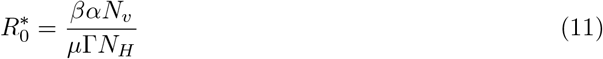

is the basic reproductive number of the temperature-independent model and Θ(Γ, *T*) encodes the effect of temperature-dependent within-host dynamics.

Two limiting cases are used repeatedly below. First, the simplified case of constant temperature with no regression, where the system is then autonomous, and the basic reproduction number can be computed exactly,

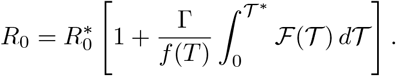

For seasonal temperature profiles that never cross the regression threshold, *R*_0_ can be approximated by replacing *f* (*T*) by its annual mean, ⟨*f* ⟩,

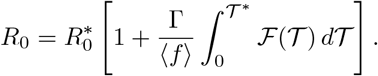

### Relating the basic reproduction number to the final epidemic size

For the autonomous version of the model, *R*_0_ is related to the final epidemic size *R*_*H*_ (∞) through

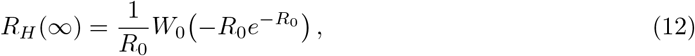

where *W*_0_ is the principal branch of the Lambert *W* function. This relation, originally derived for the classical SIR model [1] and extended to models with gamma-distributed infectious periods [53], provides an accurate final-size relation under constant temperature, when transition rates are time-independent. For time-varying temperature, Eq. (12) does not hold exactly; we use it as a benchmark to assess how informative *R*_0_ remains about final epidemic size under non-autonomous forcing.

### Simulating seasonal temperature forcing

To characterize the effect of climate on epidemic dynamics, we integrated Eq. (1)–Eq. (8) under an idealized seasonal temperature profile,

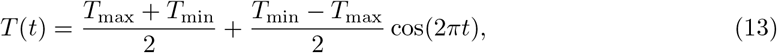

where *t* is measured in years, and *T*_min_ and *T*_max_ denote the annual minimum and maximum temperatures. This forcing reaches *T*_min_ at integer values of *t* and *T*_max_ at half-integer values of *t*, reproducing the dominant seasonal component of empirical temperature series (see Supplementary Fig. 3). The resulting dynamics for five representative temperature regimes are shown in Fig. 2.

### Assessing the effect of noise in temperature time series

To examine how short-term climatic variability modifies the dynamics obtained under purely seasonal forcing, we added stochastic fluctuations to Eq. (13). For the stochastic simulations, we integrated the model using

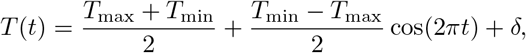

where *δ* ∼ *N*(0, *σ*^2^) is an independent Gaussian perturbation drawn at each time step

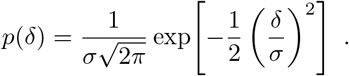

This formulation does not reproduce the autocorrelation structure of real weather variability but provides a controlled test of whether short-term fluctuations alter the epidemic outcomes obtained under smooth seasonal forcing.

As previously shown, the quantities ⟨*f* ⟩ and ⟨*g*⟩ govern the climatic contribution to invasion through *R*_0_. We therefore use the noisy temperature series to assess how variability modifies these effective progression and regression rates across climatic regimes. Using Jensen’s inequality (see Supplementary Section 7), it can be shown that variability can only increase, or leave unchanged, the mean regression rate ⟨*g*⟩, while its effect on the mean progression rate ⟨*f*⟩ depends on the shape of *f* (*T*) and the noise amplitude *σ*.

### Simulations under empirical temperature time series

We integrated the model using hourly temperature series from locations where Pierce’s disease or its causal agent, *Xylella fastidiosa* subsp. *fastidiosa* ST1, has been reported. Temperature data were obtained from the ERA5-Land dataset [54], which provides hourly temperature at approximately 10 km spatial resolution. These series were used to drive the temperature-dependent rates *γ*(*T* (*t*)) and *χ*(*T* (*t*)) and to generate the results shown in Fig. 5. Representative series and their sinusoidal fits are shown in Supplementary Fig. 3.

## Results

### Climate conditions drive disease dynamical regimes

The overall structure of the model, detailing the compartments and transitions for both hosts and vectors, is visualized in Fig. 1(a). To illustrate the mechanistic coupling between temperature and disease dynamics, we show an example of the model dynamics from *t* = 0 to *t* = 1 (in steps of Δ*t* = 0.1), with temperature following a sinusoidal pattern (see Methods). Fig. 1 (b) illustrates the time-varying nature of the physiological rates. It shows the pathogen load growth rate, *dT/dt*, as a function of time, derived from the underlying fluctuating temperature curve *T* (*t*). These time-dependent rates govern the host progression, *γ*(*T* (*t*)), and regression, *χ*(*T* (*t*)), through the infected stages.

**Figure 1.**
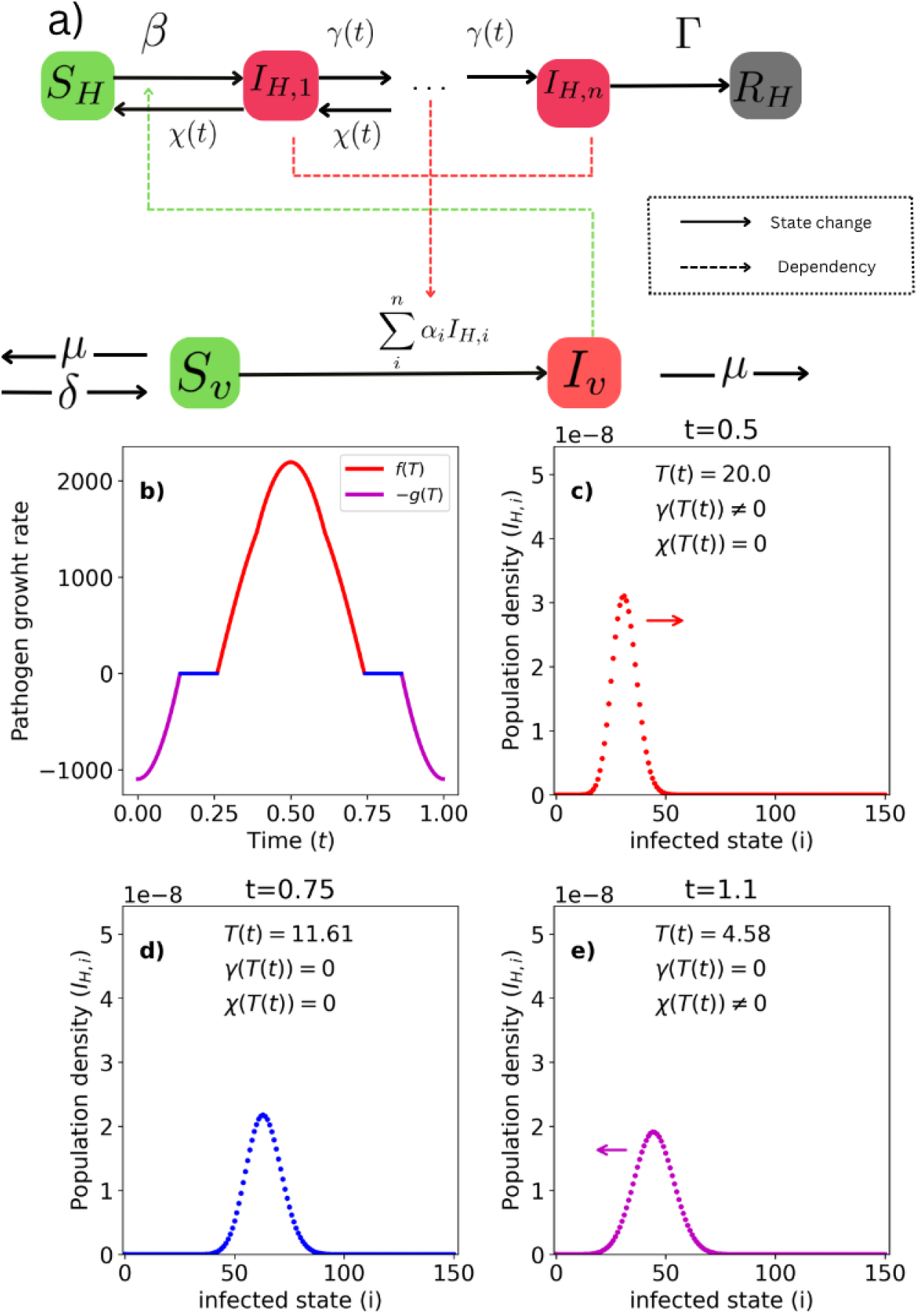
Temperature couples within-host pathogen dynamics to transmission in the stage-structured epidemic model. (a) Schematic representation of the model. Susceptible hosts (*S*_*H*_) become infected through contact with infectious vectors and progress through *n* ordered infection stages (*I*_*H*,1_, …, *I*_*H,n*_), representing increasing pathogen load and host infectiousness. Progression and regression between stages occur at temperature-dependent rates *γ*(*T* (*t*)) and *χ*(*T* (*t*)), respectively. Hosts in the final stage are chronically infected and are removed at a rate Γ. Susceptible vectors (*S*_*v*_) acquire infection from infected hosts at stage-specific rates *α*_*i*_ and remain infectious (*I*_*v*_) until death. (b) Example seasonal temperature profile and the corresponding pathogen growth and decay functions, *f* (*T* (*t*)) and *g*(*T* (*t*)). (c–e) Snapshots of the infected-host distribution across infection stages at three representative times: (c) disease progression under favorable temperatures; (d) stasis when both rates vanish; (e) regression under cold conditions. The black curve shows the stage-specific infectiousness function *F*(*Τ*_*i*_), highlighting how thermal conditions alter the overlap between within-host disease state and transmission potential.

The consequences of these fluctuating rates on the host population structure are then shown in Fig. 1 (c-e) of Fig. 1, which depict the temporal evolution of the infected host distribution (*I*_*H,i*_) across disease stages. Fig. 1 (c) captures a phase of disease progression (*t* = 0.5, *T* (*t*) = 20 °C), where temperatures favor pathogen growth (*f* (*T*) *>* 0 = *γ*(*T* (*t*)) ≠ 0). The cohort of infected hosts advances toward compartments with higher *i* values (i.e., higher pathogen load and, hence, higher infectiousness). Notably, at this early stage, the accumulated pathogen load remains insufficient to trigger significant infectiousness (black line, *F*(*T*_*i*_) ≈ 0), effectively representing a latent period. Fig. 1 (d) illustrates a state of physiological stasis (*t* = 0.75, *T* (*t*) = 11.61 °C), where the temperature falls within the range where neither growth nor decay occurs (*T*_*r*_ ≤ *T* ≤ *T*_*base*_, thus *γ*(*T* (*t*)) = *χ*(*T* (*t*)) = 0), and the host distribution remains frozen. If hosts have already reached intermediate or advanced stages, this stasis creates a window of sustained transmission potential. Fig. 1 (e) shows disease regression driven by cold temperatures (*t* = 1.1, *T* = 4.58 °C).

The activation of the decay rate (*g*(*T*) *>* 0 = *χ*(*T* (*t*)) ≠ 0) reverses the flow of infected hosts, shifting the distribution back toward earlier stages with lower pathogen loads. Consequently, the population’s overlap with the infectiousness curve decreases, thereby reducing overall transmission relative to the stasis phase.

Seasonal temperature forcing alone generates qualitatively different epidemic regimes by controlling whether infected hosts progress, stall, or regress across infection stages (Fig. 2).

**Figure 2.**
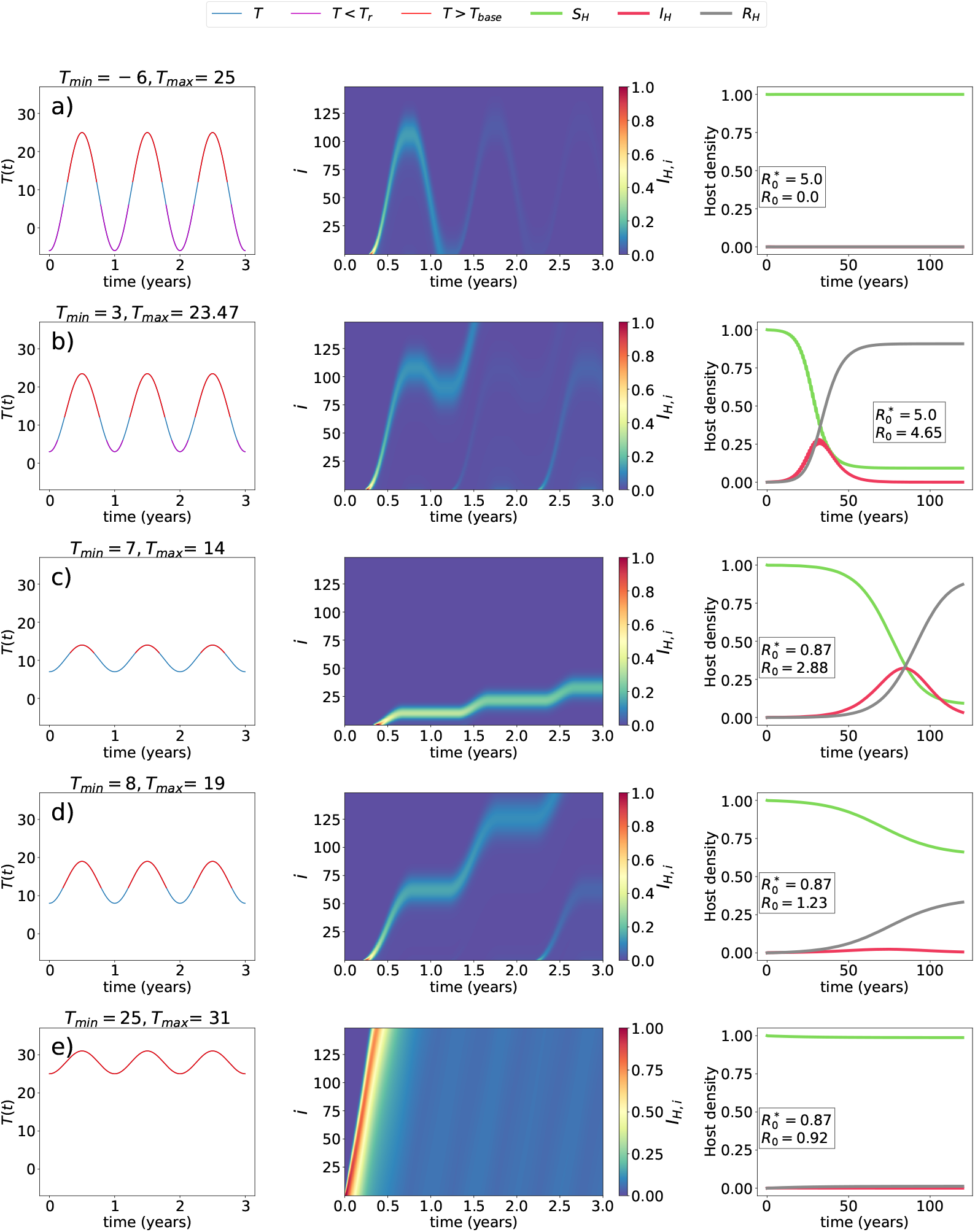
Seasonal temperature forcing alone generates qualitatively different epidemic regimes. (a–e) First column: sinusoidal seasonal temperature profiles. Red shading indicates periods of pathogen progression (*γ*(*T* (*t*)) *>* 0); blue shading, periods of thermal stasis in which neither progression nor regression occurs (*γ*(*T* (*t*)) = *χ*(*T* (*t*)) = 0); purple shading, periods of disease regression (*χ*(*T* (*t*)) *>* 0). Second column: distribution of infected hosts across infection stages, where color denotes the fraction of infected hosts in stage *i* at time *t*. Third column: corresponding host population dynamics, showing the densities of susceptible (*S*_*H*_), infected (*I*_*H*_), and removed (*R*_*H*_) hosts. Rows (a,b) illustrate that climate can suppress or permit epidemic invasion when the temperature-independent system predicts invasion 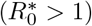. Rows (c–e) show the converse: climate can generate or suppress epidemics when the temperature-independent system predicts epidemic failure 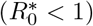.

For a fixed set of epidemiological parameters, changing the temperature profile can switch the system between epidemic suppression and sustained spread—even when the temperature-independent model would predict the opposite outcome. This is already evident for parameter sets with 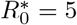. Under a colder seasonal profile (Fig. 2(a)), infected hosts progress during the warm season, but winter temperatures interrupt progression and prevent sustained transmission, so the epidemic eventually dies out. Climate can therefore suppress epidemics even when the temperature-independent system predicts invasion. Under a slightly warmer seasonal profile, however, the same within-host mechanisms permit a large epidemic outbreak to develop (Fig. 2(b)).

The converse effect emerges for parameter sets with 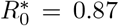, which would not produce epidemics in the absence of temperature effects. Here, cooler seasonal regimes slow progression through the infectious stages, extending the time available for transmission and allowing epidemics to develop despite 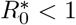 (Fig. 2(c,d)). Climate can therefore also generate epidemics when the temperature-independent system predicts failure. Under warmer regimes, however, infected hosts traverse the non-chronic infectious stages too rapidly and are removed before generating enough secondary infections to sustain spread (Fig. 2(e)).

These examples reveal that epidemic severity is not maximized by the fastest within-host progression. Temperature shapes epidemic potential through two coupled mechanisms: it determines how long infected hosts remain in the non-chronic infectious stages that strongly contribute to transmission, and whether regression becomes possible under unfavorable conditions. Slightly less favorable thermal conditions can generate larger epidemics than warmer ones by prolonging time in highly infectious stages, whereas colder conditions suppress spread by halting progression or inducing regression. This balance between transmission opportunity, progression speed, and recovery provides the biological intuition for the analytical results that follow.

### How temperature shapes epidemic invasion

The contrasting epidemic outcomes described above can be understood analytically through the model’s basic reproduction number (see Methods). Under seasonal temperature forcing, this quantity decomposes Eq. (10) into a temperature-independent component 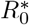 and a temperature-dependent modulation factor Θ(Γ, *T*) that captures the combined effect of mean progression rate ⟨*f*⟩ and, when temperatures fall sufficiently low, mean regression rate ⟨*g*⟩.

The limiting case of constant temperature makes this mechanism particularly transparent. When pathogen growth is possible, and regression is absent, the general expression for *R*_0_ Eq. (10) reduces to

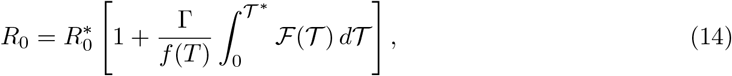

which is the exact basic reproduction number of the autonomous system (see Methods).

Across both constant and seasonal temperature profiles, simulation outcomes collapse onto a narrow relationship between the final epidemic size, *R*_*H*_ (∞), and the corresponding temperature-dependent reproduction number (Fig. 3 a-b). The collapse is tightest under constant temperature (Fig. 3 a) and somewhat broader under time-varying forcing (Fig. 3 b), where the residual scatter concentrates near the epidemic threshold. As shown below, this is precisely where the averaging that underlies our analytical *R*_0_ is expected to lose accuracy, because the temporal ordering of warm and cold periods—discarded by the annual average—begins to matter. This shows that temperature primarily shapes epidemic invasion by modifying the number of secondary infections generated during an infection.

The climate-dependent contribution to the constant-temperature expression Eq. (14) is inversely proportional to the within-host pathogen growth rate, *f* (*T*). Temperature, therefore, does not enhance epidemic invasion simply by accelerating pathogen growth. Instead, it changes the duration of the non-chronic infectious stages in infected hosts, which can substantially contribute to transmission. When *f* (*T*) is small, infected hosts progress more slowly through those stages and accumulate more opportunities to infect vectors, increasing *R*_0_. When *f* (*T*) is large, those stages are traversed rapidly, and their contribution becomes negligible. In the limit of infinitely fast within-host progression, the thermal influence on the basic reproductive number vanishes,

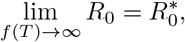

and the model reduces to its temperature-independent form.

This limiting behavior explains the counterintuitive regimes identified in Fig. 2. For parameter sets with 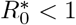, temperatures near the within-host optimum maximize *f* (*T*) and drive *R*_0_ back toward 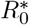, so that invasion fails even though pathogen growth within hosts is fastest. By contrast, mildly suboptimal temperatures slow progression, prolong the time spent in infectious stages, and can raise *R*_0_ *>* 1, generating large outbreaks. Conversely, when 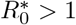, cold seasonal conditions can suppress epidemics by interrupting progression or, if temperatures drop below the regression threshold, by allowing infected hosts to move backward through the infection chain and recover, thereby reducing the effective reproductive number to *R*_0_ *<* 1.

The same interpretation carries over to seasonal forcing. When temperatures never fall below the regression threshold (*T*_*min*_ *> T*_*r*_), the approximation for *R*_0_ takes the same form as the constant-temperature result Eq. (14) after replacing *f* (*T*) by its annual mean, ⟨*f*⟩ : seasonal fluctuations matter mainly through their effect on average progression speed, and the same residence-time mechanism explains why slower seasonal disease development can yield larger epidemics. When temperatures do cross the regression threshold (*T*_*min*_ *< T*_*r*_), epidemic invasion depends on the balance between average progression and average regression, captured by the ratio ⟨*g*⟩ */* ⟨*f*⟩. Increasing ⟨*g*⟩ decreases epidemic invasion by promoting recoveries, whereas decreasing net progression increases the time spent in infectious stages and can enhance transmission. Near the transition ⟨*g*⟩ */* ⟨*f*⟩ ≈ 1, these effects nearly compensate, and the temporal ordering of warm and cold periods becomes important, reducing the accuracy of the averaged approximation. Even in that regime, however, the analytical result retains its biological interpretation: epidemic invasion is determined by the competition between transmission accumulated during slow progression and recovery induced by unfavorable temperatures—consistent with the increased scatter of *R*_*H*_ (∞) about the final-size relation in this regime (Fig. 3b).

**Figure 3.**
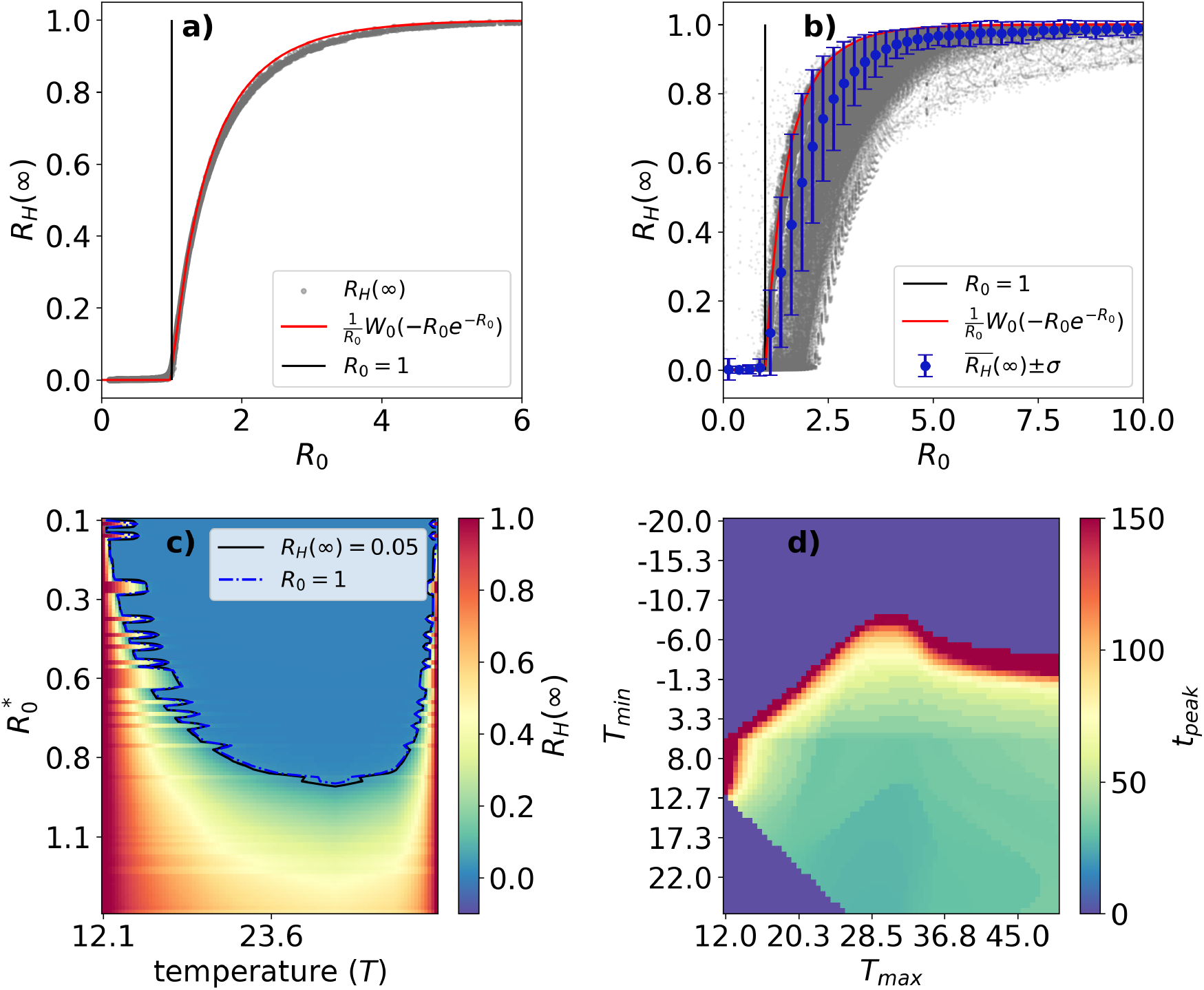
The approximate temperature-dependent reproduction number is strongly predictive of final epidemic size. (a) Gray dots represent the final epidemic size obtained from simulations under constant temperature forcing. The solid red curve shows the classical final-size Relation 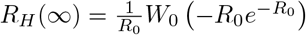, where *W*_0_ is the principal branch of the Lambert *W* function, and the epidemic threshold, *R*_0_ = 1, is indicated by the vertical line. (b) Final epidemic size from simulations under time-varying temperature forcing plotted against the corresponding approximate temperature-dependent reproduction number; gray points show individual simulations, the red curve is the classical final-size relation between *R*_0_ and the final epidemic size, the black vertical line indicates *R*_0_ = 1, and blue symbols show the mean ± standard deviation of *R*_*H*_ (∞) within bins of *R*_0_. (c) Final epidemic size under constant temperature across the (*T*, 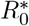) parameter space; the blue dashed curve marks *R*_0_ = 1 and the black curve marks *R*_*H*_ (∞) = 0.05. (d) Time to epidemic peak across the (*T*_min_, *T*_max_) climate space, illustrating how thermal regime shapes not only invasion probability but epidemic dynamics.

Overall, the analytical expressions show that temperature shapes epidemic invasion not simply by speeding up or slowing down pathogen growth within hosts, but through its combined effects on time spent in infectious stages and on the possibility of regression and recovery. This explains both climate-driven suppression of epidemics and emergence, and why the temperatures most favorable for within-host pathogen growth are not necessarily those that maximize epidemic spread.

### The effect of climate variability

We examined whether short-term variability around the seasonal climate cycle modifies the invasion patterns described above. Adding stochastic fluctuations to seasonally forced temperatures preserves the same underlying mechanism, but alters the effective progression and regression experienced by infected hosts (Fig. 4a–c). Variability can increase regression by intermittently pushing temperatures below the regression threshold, while it modifies progression asymmetrically depending on the underlying seasonal regime. When the baseline seasonal profile keeps temperatures near the within-host optimum for pathogen growth, fluctuations shift portions of the thermal trajectory toward less favorable values, reducing mean progression. Under more suboptimal regimes, variability can occasionally move temperatures toward the optimum and increase mean progression. As a result, the climatic contribution to epidemic invasion, measured by *R*_0_, decreases across most of the (*T*_*min*_, *T*_*max*_) climate space and increases only in a relatively small region of warm seasonal profiles (Fig. 4c).

**Figure 4.**
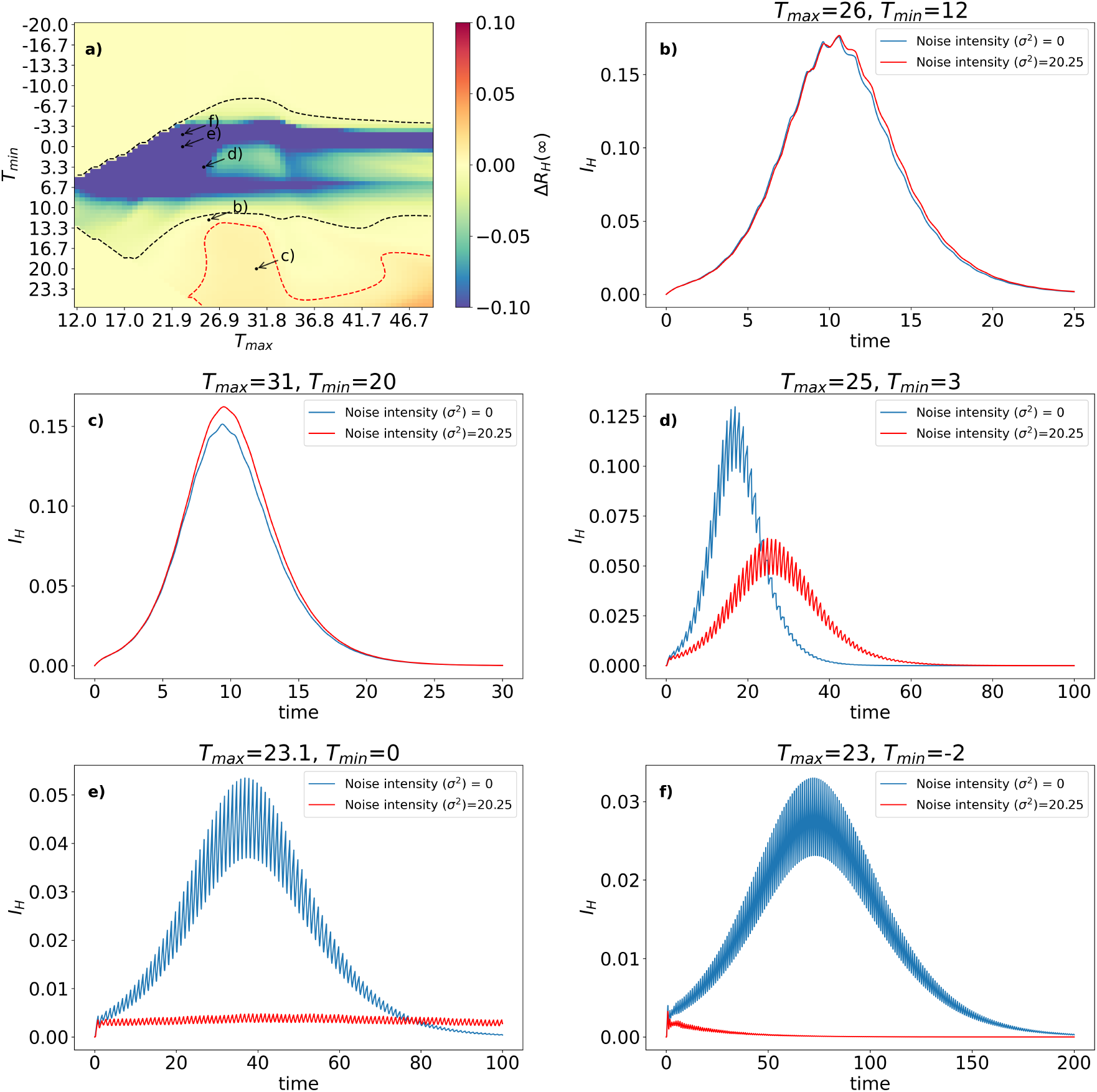
Short-term climatic variability mainly suppresses epidemic invasion near thermal thresholds. (a) Noise-induced changes in final epidemic size, Δ*R*_*H*_ (∞), across the (*T*_*min*_, *T*_*max*_) climate space; solid black and dashed red contours delimit regions where variability decreases or increases *R*_*H*_ (∞), respectively. (b–f) Representative epidemic trajectories for the seasonal profiles marked in panel (a), illustrating cases in which variability has a negligible effect, slightly enhances epidemic size, weakens and delays epidemics, or suppresses them altogether. Insets show the corresponding temperature series with and without noise.

These changes translate directly into epidemic outcomes. Across most seasonal temperature profiles, adding variability either leaves final epidemic size nearly unchanged or reduces it, with the strongest reductions occurring near the thermal boundary between epidemic invasion and epidemic failure (Fig. 4a). In these near-threshold climates, epidemic spread depends on a delicate balance between slow progression and limited recovery, so short-term fluctuations can qualitatively alter the dynamics. Epidemics that occur under smooth sinusoidal forcing may become slower and smaller (Fig. 4d), collapse into long-lasting low-level infections with almost no host mortality (Fig. 4e), or disappear altogether (Fig. 4f).

By contrast, in warm seasonal regimes far from the regression threshold, the effect of variability is weak and can even slightly increase epidemic size (Fig. 4b,c). In this part of the climate space, fluctuations do not cause substantial regression; their main consequence is to transiently displace temperatures away from the pathogen growth optimum, slowing progression through the non-chronic infectious stages and modestly prolonging the window available for transmission. This enhancement effect, however, is small compared with the suppressive effect near threshold climates.

Overall, climate variability does not create qualitatively new epidemic regimes. Rather, it modulates the same trade-off between progression speed, time in infectious stages, and recovery identified above. Its dominant effect is to reduce epidemic invasion by intermittently creating unfavorable thermal conditions, particularly in climates already near the transition between epidemic spread and failure. This makes epidemic outcomes more sensitive to short-term fluctuations and provides a natural bridge to the empirical temperature series analyzed next.

### Empirical temperature series reproduce contrasting epidemic behaviors across invaded regions

We applied the model to empirical temperature series from six locations where Pierce’s disease has been reported: Sierra de Gata (Extremadura, Spain), Pegarinhos (Bragança, Portugal), Santa Maria del Camí (Mallorca, Spain), Bari (Apulia, Italy), Castelo de Marvão (Portalegre, Portugal), and Fundão (Castelo Branco, Portugal). (Fig. 5). In all six cases, the model predicts epidemic invasion, consistent with the corresponding temperature-dependent reproduction numbers exceeding one. However, the simulated dynamics differ markedly among sites, showing that empirical thermal regimes can generate distinct epidemic trajectories even when all locations remain climatically suitable for invasion.

**Figure 5.**
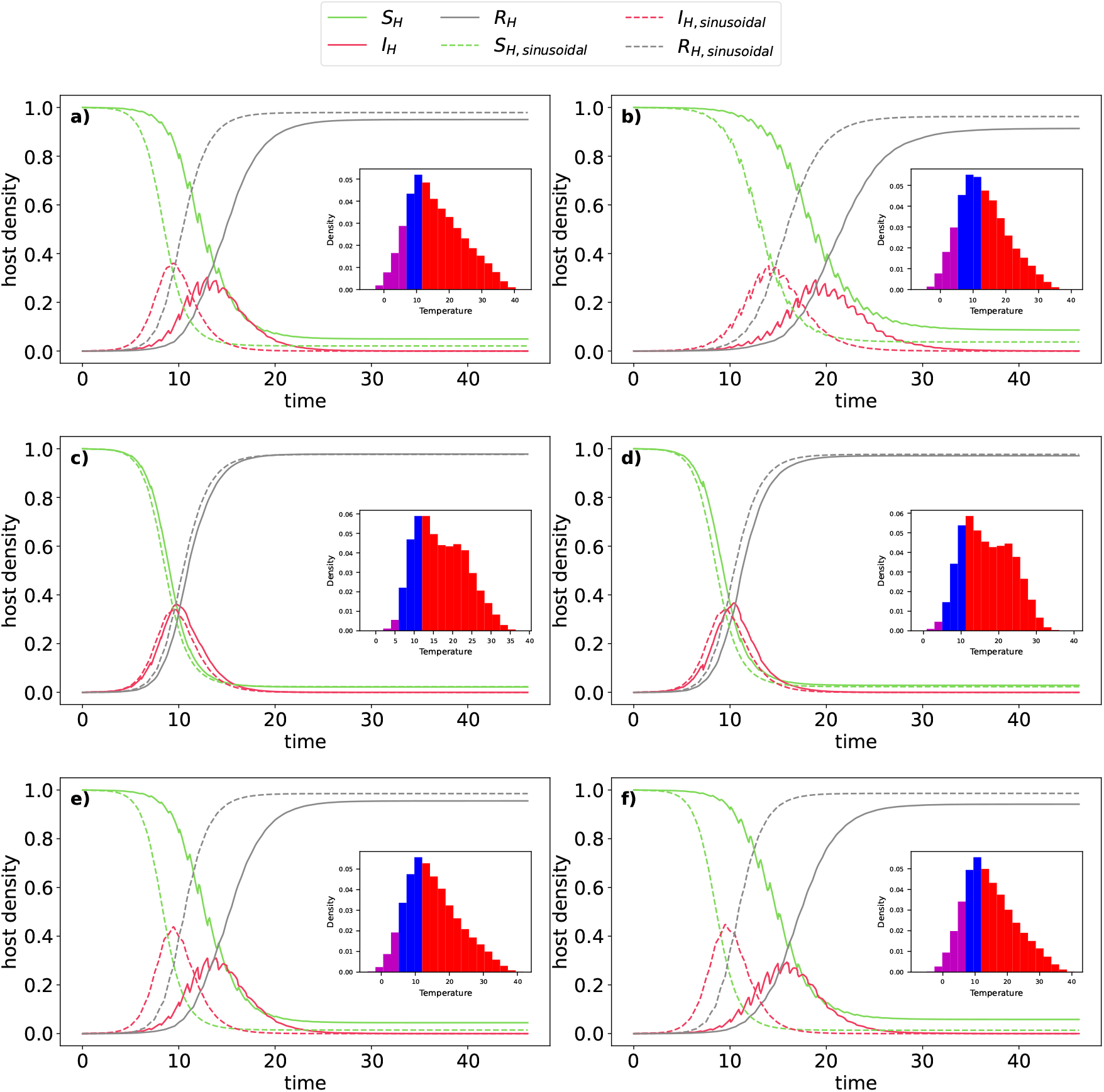
Empirical temperature series reproduce contrasting epidemic behaviors across invaded regions. Host dynamics under the empirical temperature series (solid lines) and the corresponding fitted sinusoidal temperature profile (dashed lines), showing susceptible (*S*_*H*_), infected (*I*_*H*_), and removed (*R*_*H*_) individuals for six locations where Pierce’s disease has been reported: (a) Sierra de Gata (Extremadura, Spain), (b) Pegarinhos (Bragança, Portugal), (c) Santa Maria del Camí (Mallorca, Spain), (d) Bari (Apulia, Italy), (e) Castelo de Marvão (Portalegre, Portugal), and (f) Fundão (Castelo Branco, Portugal). Insets show the histograms of the empirical temperature series. Together, these examples illustrate how empirical thermal regimes can generate markedly different epidemic dynamics, even when all sites remain climatically suitable for invasion.

Two broad dynamical patterns emerge. In Apulia and Mallorca, epidemics develop comparatively rapidly with little evidence of seasonal regression: infected hosts accumulate pathogen load continuously across the year, and epidemic growth is driven mainly by progression to advanced infectious stages. In the remaining sites, by contrast, epidemics are substantially slower and display clear seasonal modulation, with recurrent annual declines in infection consistent with partial recovery (regression) and reinfection. These differences are biologically meaningful: they indicate that some invaded regions are expected to support direct progression toward severe infection, while in others the pathogen may persist through slower, seasonally interrupted trajectories—a distinction with direct implications for symptom timing, the duration of the infectious window, and the pace at which vineyards are affected.

These patterns reflect differences in winter thermal regimes across the six sites. In Apulia and Mallorca, the absence of sufficiently cold winter nights means that no winter-curing effect occurs and disease progression is uninterrupted throughout the year. At the remaining sites, a more continental influence lowers winter temperatures below the regression threshold, partially offsetting progression; however, the net annual balance still favors overall disease advance, consistent with the model’s prediction of invasion at both sites. This interpretation is confirmed by the geography of the selected locations: the Apulia and Mallorca sites are coastal, while the remaining sites are situated further inland Fig. 5.

This contrast is organized by the same thermal metrics identified in the previous sections. Sites with negligible regression are characterized by large net annual progression, so infected hosts approach the chronic stage within roughly one growing season, and epidemic peaks occur earlier. Sites with stronger regression show much smaller net annual increase in infection, delaying the accumulation of highly infectious hosts and shifting the epidemic peak to substantially later times. The balance between progression and regression, therefore, determines not only whether local epidemics are fast or slow, but whether infection advances steadily or through repeated seasonal setbacks—in idealized and empirical climates alike.

The empirical series also corroborate the conclusions of the variability analysis. Relative to their fitted seasonal baselines, short-term variability tends to delay epidemic development and reduce final epidemic size, with the strongest effects at the sites where seasonal regression is already important. In the warmer sites with little regression, deviations from the seasonal cycle have modest epidemiological consequences. In cooler sites, variability amplifies seasonal interruptions in disease progression, resulting in substantially slower and smaller epidemics. Empirical temperature series, therefore, do not introduce qualitatively new behaviors; they combine the mechanisms identified for seasonal forcing and short-term variability, and confirm that both are relevant for understanding how epidemic dynamics may differ across invaded landscapes.

## Discussion

By explicitly linking temperature-dependent within-host pathogen progression to vector-mediated transmission, our framework addresses a central challenge in plant disease epidemiology: representing time-varying infectiousness arising from physiological processes unfolding within infected hosts [21, 22]. Most epidemic models treat infectiousness as fixed or impose environmental forcing directly on transmission coefficients—a practical choice, but one that weakens the connection between controlled physiological measurements and epidemic dynamics under fluctuating conditions. Here, we show that mechanistic coupling between pathogen load dynamics and transmission can bridge that gap: climate affects epidemic behavior not only by altering invasion likelihood but also by reshaping the duration of infectious stages, the possibility of regression, and the accumulation of transmission over time. This provides a tractable route for scaling experimentally measurable within-host processes into population-level epidemic dynamics under non-stationary climatic forcing. Our results extend recent progress in modeling *Xylella fastidiosa* diseases under environmental forcing. Previous climate-driven models have shown that temperature explains current disease distributions and future epidemic risk [32–34], but focused primarily on potential establishment. Our framework shifts the question of focus from whether invasion occurs to how epidemics unfold once established, a distinction with direct consequences for forecasting and management. This is consistent with recent work emphasizing that climate sensitivity in infectious diseases emerges from nonlinear effects of environmental forcing on pathogen, host, and vector traits rather than from simple monotonic responses [55, 56]. Field-based studies have likewise shown that warmer conditions and milder winters can increase *X. fastidiosa* outbreak risk by affecting vector infection prevalence and climatic suitability [57], and recent work has further highlighted the importance of environmental and ecological context for transmission, spread, and management across *Xylella* pathosystems [58–60].

A central mechanistic result is that epidemic spread is not maximized by the fastest within-host pathogen growth. The temperature-dependent component of the basic reproduction number *R*_0_ reflects a trade-off between progression speed and the time infected hosts spend in non-chronic infectious stages that contribute to transmission. Temperatures mildly suboptimal for pathogen development can produce larger epidemics than temperatures near the within-host optimum, because they prolong residence times in infectious stages and increase cumulative transmission: a slow-growth paradox. This provides a concrete example of a point increasingly recognized in infectious disease theory: transient within-host dynamics can have major population-level consequences and should not be treated as decoupled from between-host transmission [61]. In plant disease systems, where temperature affects pathogen multiplication, symptom expression, and recovery, explicitly linking these processes to transmission may reveal dynamical regimes that remain hidden in models that force transmission parameters in aggregate.

A second important implication concerns the dual role of seasonal climate in shaping invasion. Seasonal forcing influences epidemic dynamics through two coupled pathways: modulating the rate of progression and inducing regression. As a result, climate can suppress epidemics even when a temperature-independent model predicts invasion, and permit them when such a model predicts failure. These dynamics are especially sensitive near the boundary between progression-dominated and regression-dominated regimes, where small thermal shifts determine whether infected hosts accumulate sufficient pathogen load to sustain transmission. These findings complement recent climate-risk studies of Pierce’s disease that have highlighted strong sensitivity of epidemic potential to fine-scale thermal conditions [33], and fit within a broader literature showing that climate effects on infectious diseases are often delayed, threshold-dependent, and mechanistically structured across multiple biological scales [56].

The empirical temperature series reinforces this picture while adding an important nuance: climatically suitable invaded regions need not support similar epidemic trajectories. All six sites remain permissive for invasion, yet differ strongly in dynamics, consistent with evidence that *Xylella fastidiosa* risk varies markedly across regions with distinct thermal regimes [33, 57]. In Apulia and Mallorca—coastal sites subject to strong maritime influence and mild winters—the absence of a winter-curing effect allows uninterrupted disease progression, leading to rapid advancement to later infectious stages. In the remaining sites, a more continental thermal regime brings winter temperatures below the regression threshold, resulting in slower, seasonally interrupted epidemics in which infection persists for extended periods before substantial host mortality accumulates. The net annual balance still favors overall disease advance in both regimes, but the trajectory is qualitatively different—a distinction consistent with experimental and field evidence that milder winters favor pathogen persistence and hotter growing seasons limit recovery [43, 57, 62].

These differences carry both predictive and practical implications. On the predictive side, the specific Apulian location where Pierce’s disease has been detected lies outside the Salento Peninsula— epicenter of the severe olive-tree epidemic caused by *Xylella fastidiosa* subsp. *pauca*—and, given its stronger maritime influence and mild winter nights, would be expected to behave epidemiologically more like Mallorca than like the continental Apulian interior, with faster progression and limited seasonal regression. More broadly, local thermal regimes shape not only outbreak probability but also symptom timing, the persistence of infected but non-terminal hosts, and the duration of the window during which vectors can acquire and redistribute the pathogen [57, 63]. On the practical side, this means that the absence of a rapid, visible decline should not be interpreted as evidence of low epidemiological relevance in cooler, invaded regions: transmission may be ongoing and substantial even when symptom expression remains limited, with direct consequences for the design of surveillance strategies and the timing of management interventions.

Our analysis of climate variability reinforces and sharpens this point. Short-term fluctuations around the seasonal cycle do not create fundamentally new epidemic regimes, but they can strongly modulate the balance among progression, infectious residence time, and recovery. Their effects are largest near thermal thresholds, where intermittent excursions below the regression boundary can transform an epidemic predicted under smooth seasonal forcing into a substantially smaller outbreak or suppress it entirely. In already warm regimes, variability has weaker effects and can occasionally increase epidemic size slightly by transiently slowing progression away from the thermal optimum. This asymmetry is consistent with recent arguments that nonlinear environmental trait responses—including delayed responses of vectors—can substantially alter disease forecasts and should be incorporated into mechanistic models wherever possible [56, 64].

Several limitations of the present framework merit consideration, two of which bound the regime in which the slow-growth mechanism operates. First, the model assumes that infected hosts do not undergo temperature-independent clearance or removal before reaching the chronic stage. This approximation is appropriate for Pierce’s disease, for which documented recovery is primarily associated with cold exposure and is represented here through temperature-dependent regression. In systems with appreciable background mortality or loss of pre-chronic hosts, an additional exit rate would compete with thermal progression and limit the time available for transmission; the slow-growth mechanism identified here therefore applies most directly when such temperature-independent losses are negligible. Second, temperature acts in our model exclusively through within-host pathogen dynamics, with vector abundance, phenology, and competence treated as climate-independent. This was a deliberate simplification to isolate the epidemiological consequences of temperature-driven disease progression, but it likely underestimates the full range of climate effects relevant to Pierce’s disease and other vector-borne plant pathogens. Indeed, climate can act on pathogens and vectors in opposing directions [65], and we have shown for Pierce’s disease in Europe that warming can simultaneously expand pathogen suitability while contracting the climatic range of *Philaenus spumarius* [34]. Coupling climate-dependent vector demography and behavior to the present within-host framework is therefore an important direction for future work.

Beyond these structural assumptions, several caveats concern the model’s scope and empirical application. External interventions such as roguing or pruning of infected tissue are not represented, as they act on the host population rather than on intrinsic within-host dynamics; incorporating them as removal terms is straightforward within the present structure. The stochastic temperature fluctuations considered are temporally uncorrelated and therefore do not fully capture the persistence structure of real weather variability or the epidemiological consequences of extreme events. Finally, the temperature series presented are illustrative rather than fitted reconstructions of observed outbreaks, and the framework has not yet been calibrated against field incidence data. Addressing these points—incorporating delayed trait plasticity, landscape heterogeneity, and field-calibrated host–vector interactions—would be natural extensions of the present framework, and would align it with emerging integrative approaches across *Xylella* pathosystems [60].

Despite these possible extensions, the present results point to a specific and transferable conclusion: stage-structured infectiousness under thermal forcing can generate a slow-growth paradox. Whenever temperature jointly controls pathogen development, symptom progression, and host recovery, models that do not resolve within-host dynamics may fail not only quantitatively but also qualitatively, misidentifying the thermal regimes that pose the greatest epidemic threat.

## Data and code availability

No new experimental data were generated in this study. The model was parameterized using previously published experimental data on temperature-dependent *Xylella fastidiosa* growth, symptom progression, and recovery in grapevine [32]. ERA5-Land temperature data were obtained from the Copernicus Climate Change Service Climate Data Store [54]. The source code used to implement the model, compute the temperature-dependent progression and regression rates, reproduce all numerical simulations, and generate the figures is publicly available through GitHub [66] and archived with a permanent DOI in Zenodo [67]. The archived repository includes the processed temperature series, parameter files, analysis scripts, and documentation needed to reproduce the results.

## Acknowledgements

We thank Eduardo Moralejo and Frederic Bartumeus for feedback in previous versions of this work. We acknowledge support by grants PID2024-156062OB-I00 (CHANGE-ME), funded by the Spanish Ministry of Science and Innovation MICIU/AEI/10.13039/501100011033 and by ERDF, EU; CEX2021-001164-M (María de Maeztu Program for Units of Excellence in R&D) funded by MICIU/AEI/10.13039/501100011033; MEPRO 4832/2024 funded by Conselleria d’Agricultura del Govern de les Illes Balears. AGR acknowledges financial support from grant JDC2024-053275-I, funded by MICIU/AEI/10.13039/501100011033 and FSE+.

## Author contributions statement

MAM and AGR conceived the study. JCRC, MAM, and AGR designed the theoretical framework. JCRC developed and implemented the model. JCRC performed the numerical simulations and formal analyses, with input from AGR. JCRC, MAM, and AGR wrote the manuscript. AGR and MAM supervised the study. MAM obtained the funding.

