## Supplementary Information for "A mechanistic framework linking within-host pathogen progression to vector-mediated transmission under climate forcing"

*Note on notation.* The thermal integral for growth is denoted MGDD in the original study [1]. We refer to it as GDD throughout this manuscript for notational simplicity. Analogously, the Killing Degree Days (KDD) in this study were referred to as CDD in [1]. Furthermore, references to equations as a single number, without a dot, refer to equations in the Main Article, while references with two numbers separated by a dot, say (1.1), refer to equations in this SI.

#### 1 Parameterization for Pierce's disease

We parameterize the model using experimental data for Pierce's disease of grapevine, caused by *Xylella fastidiosa* [1, 2]. The temperature-dependent bacterial growth function, $f(T)$, is derived from experimental data on pathogen growth [2] and is defined by

$$f(T) = \begin{cases} 0 & \text{if} \quad T < T_{\text{base}} \\ m_1 \cdot T + b_1 & \text{if} \quad T_{\text{base}} \leq T < T_1 \\ m_2 \cdot T + b_2 & \text{if} \quad T_1 \leq T < T_{\text{opt}} \\ m_3 \cdot T + b_3 & \text{if} \quad T_{\text{opt}} \leq T < T_2 \\ m_4 \cdot T + b_4 & \text{if} \quad T_2 \leq T < T_{\max} \\ 0 & \text{if} \quad T \geq T_{\max} \end{cases}$$

where $T_{\text{base}} = 12\,°\text{C}$, $T_1 = 18$, $T_{\text{opt}} = 28\,°\text{C}$, $T_2 = 32$ and $T_{\max} = 35\,°\text{C}$; the slopes are $m_1 = 0.66$, $m_2 = 1$, $m_3 = -1.25$ and $m_4 = -3$ and the intercepts are $b_1 = -8$, $b_2 = -14$, $b_3 = 4$ and $b_4 = 105$.

The thermal integral for growth is then computed as

$$\text{GDD}(t) = \int_{t_0}^{t} f(T(t'))\, dt'.$$

No direct experimental data link the bacterial death rate to temperature. However, cold-driven host recovery (winter curing) has been documented and related to cold accumulation [1, 3, 4]. Following [1], we model pathogen decay through a cold accumulation function,

$$\text{KDD}(t) = \int_{t_0}^{t} g(T(t'))\, dt', \qquad g(T) = \max(T_r - T,\ 0),$$

where $T_r = 6\,^\circ\mathrm{C}$ is the regression threshold. Together, $f(T)$ and $g(T)$ define three thermal regimes: pathogen growth occurs when $T > T_{\text{base}}$; pathogen decay occurs when $T < T_r$; and pathogen load remains constant when $T_r \le T \le T_{\text{base}}$, provided the temperature continuously remains within this range.

Notably, $f(T)$ exhibits an optimum at $T = T_{\text{opt}}$, the temperature for maximal bacterial growth, while $g(T)$ models a rate of decay that increases unboundedly as temperature decreases, suggesting rapid bacterial death at extremely low temperatures.

To link pathogen load to host infectiousness, we use experimental inoculation data in which symptomatic leaf count—a proxy for pathogen load—was tracked after inoculation alongside ambient temperature [1]. Because the experiments were conducted under warm conditions throughout, no cold accumulation occurred (KDD = 0), and all symptom development was attributable to GDD alone. A threshold of five symptomatic leaves was used to define the transition to chronic infection. The resulting probability of an infected host transitioning to the chronic state is described by the sigmoidal function

$$\mathcal{F}(GDD) = \frac{1}{1 + e^{-0.012(GDD-975)}} \ ,$$

To link within-host pathogen load to between-host transmission, we assume that the probability of a susceptible vector acquiring infection while feeding on a host in stage $i$ is proportional to the host's symptomatic load at that stage, giving the stage-specific acquisition rate,

$$\alpha_i = \alpha \mathcal{F}(\mathcal{T}_i),$$

where $\alpha$ is the acquisition rate at the chronic stage.

To derive the temperature-dependent transition rates between compartments, we define $\mathcal{T}^*$ as the value at which $\mathcal{F}(\mathcal{T}^*) = 0.99$, representing the bacterial load required for a host to become chronically infected with 99% probability. The interval $[0, \mathcal{T}^*]$ is divided into $n$ equal sub-intervals of width $\Delta\mathcal{T} = \mathcal{T}^*/n$, each corresponding to one infection stage. The progression and regression rates are then

$$\gamma(T(t)) = \frac{f(T(t))}{\Delta\mathcal{T}}. \tag{1.1}$$

$$\chi(T(t)) = \frac{g(T(t))}{\Delta\mathcal{T}}. \tag{1.2}$$

### 2 Full derivation of $R_{0,\text{av}}$

Replacing $\gamma(T(t))$ and $\chi(T(t))$ by their annual averages

$$\bar{\gamma} = \int_0^1 \gamma(T(t))\,dt, \qquad \bar{\chi} = \int_0^1 \chi(T(t))\,dt,$$

an analytical expression can be obtained for the Jacobian of the infected subsystem evaluated at the disease-free equilibrium, corresponding to the first-order Magnus approximation [5],

$$J\big|_{DFE} = \begin{pmatrix} -\mu & \alpha_1 N_v/N_H & \alpha_2 N_v/N_H & \alpha_3 N_v/N_H & \dots & \alpha_{n-1} N_v/N_H & \alpha N_v/N_H \\ \beta & -(\bar{\gamma}+\bar{\chi}) & \bar{\chi} & 0 & \dots & 0 & 0 \\ 0 & \bar{\gamma} & -(\bar{\gamma}+\bar{\chi}) & \bar{\chi} & \dots & 0 & 0 \\ \vdots & \vdots & \vdots & \ddots & \vdots & \vdots & \vdots \\ 0 & 0 & 0 & 0 & \dots & -(\bar{\gamma}+\bar{\chi}) & 0 \\ 0 & 0 & 0 & 0 & \dots & \bar{\gamma} & -\Gamma \end{pmatrix}.$$

Applying the standard next-generation decomposition $J = F - V$ , where $F$ contains new infection terms and $V$ contains all other transitions,

$$F = \begin{pmatrix} 0 & 0 & 0 & 0 & \dots & 0 & 0 \\ \beta & 0 & 0 & 0 & \dots & 0 & 0 \\ 0 & 0 & 0 & 0 & \dots & 0 & 0 \\ \vdots & \vdots & \vdots & \ddots & \vdots & \vdots & \vdots \\ 0 & 0 & 0 & 0 & \dots & 0 & 0 \\ 0 & 0 & 0 & 0 & \dots & 0 & 0 \end{pmatrix},$$

$$V = \begin{pmatrix} \mu & -\alpha_1 N_v/N_H & -\alpha_2 N_v/N_H & -\alpha_3 N_v/N_H & \dots & -\alpha_{n-1} N_v/N_H & -\alpha N_v/N_H \\ 0 & \bar{\gamma}+\bar{\chi} & -\bar{\chi} & 0 & \dots & 0 & 0 \\ 0 & -\bar{\gamma} & \bar{\gamma}+\bar{\chi} & -\bar{\chi} & \dots & 0 & 0 \\ \vdots & \vdots & \vdots & \ddots & \vdots & \vdots & \vdots \\ 0 & 0 & 0 & 0 & \dots & \bar{\gamma}+\bar{\chi} & 0 \\ 0 & 0 & 0 & 0 & \dots & -\bar{\gamma} & \Gamma \end{pmatrix}.$$

The next-generation matrix is $K = FV^{-1}$, and $R_0 = \rho(K)$, where $\rho(\cdot)$ denotes the spectral radius. The dominant eigenvalue is

$$R_{0,\mathrm{av}} = \frac{\beta}{\mu\Gamma}\frac{N_v}{N_H}\frac{1}{\sum_{j=0}^{n-1}\bar{\chi}^j\bar{\gamma}^{(n-1-j)}}\left[\Gamma\sum_{i=0}^{n-1}\bar{\gamma}^{n-2}\alpha_i + \Gamma\bar{\gamma}^{n-1}\alpha + \sum_{i=1}^{n-2}\alpha_i\sum_{k=i-1}^{n-3}\bar{\chi}^{n-2-k}\bar{\gamma}^k\right],$$

which can be explicitly written as a function of the pathogen thermal niche as

$$R_{0,\mathrm{av}} = R_0^*\left[\frac{\Gamma}{\bar{f}}\left\{\int_0^{\mathcal{T}^*}\mathcal{F}(\mathcal{T})\,d\mathcal{T} + \sum_{i=1}^{n-2}\mathcal{F}(\mathcal{T}_i)\,\Delta\mathcal{T}\sum_{k=i-1}^{n-3}\left(\frac{\bar{g}}{\bar{f}}\right)^{n-2-k}\right\}+1\right] \times \left(1-\frac{\bar{g}}{\bar{f}}\right)\Bigg/\left(1-\left(\frac{\bar{g}}{\bar{f}}\right)^{n-1}\right) \tag{2.1}$$

where

$$R_0^* = \frac{\beta\alpha N_v}{\mu\Gamma N_H} \tag{2.2}$$

is the basic reproduction number of the temperature-independent model, and $\bar{f}$ and $\bar{g}$ are the annual averages of the temperature-dependent progression and regression functions defined in Eq. (1.1)-Eq. (1.2). In deriving Eq. (2.1), we used the large-$n$ approximation

$$\sum_{i=0}^{n-1}\frac{\alpha_i}{\bar{\gamma}} \approx \alpha\int_0^{\mathcal{T}^*}\mathcal{F}(\mathcal{T})\,d\mathcal{T}\,.$$

For convenience, we define

$$\mathcal{I} = \int_0^{\mathcal{T}^*}\mathcal{F}(\mathcal{T})\,d\mathcal{T}\,,$$

$$\mathcal{I}_r(T) = \sum_{i=1}^{n-2}\mathcal{F}(\mathcal{T}_i)\,\Delta\mathcal{T}\sum_{k=i-1}^{n-3}\left(\frac{\bar{g}}{\bar{f}}\right)^{n-2-k},$$

$$\mathcal{C}(T) = \left(1-\frac{\bar{g}}{\bar{f}}\right)\Bigg/\left(1-\left(\frac{\bar{g}}{\bar{f}}\right)^{n-1}\right)\,.$$

Recall that $\bar{\gamma}$ and $\bar{\chi}$ are the annual averages of the temperature-dependent progression and regression functions defined in Eqs. (1.1) and (1.2), hence the temperature dependence of $\mathcal{I}_r(T)$ and $\mathcal{C}(T)$.

With these definitions,

$$\Theta(\Gamma, T) = \left( \frac{\Gamma}{\bar{f}} \left[ \mathcal{I} + \mathcal{I}_r(T) \right] + 1 \right) \mathcal{C}(T),$$

so that

$$R_0 \approx R_0^* \, \Theta(\Gamma, T).$$

For constant temperature, with $g(T) = 0$ (no regression) we have $\bar{\chi} = 0$, $\mathcal{I}_r = 0$, and $\mathcal{C}(T) = 1$, so Eq. (2.1) reduces to

$$R_0 = R_0^* \left[ 1 + \frac{\Gamma}{f(T)} \mathcal{I} \right].$$

Because the system is then autonomous, this is the exact basic reproduction number. For seasonal temperature profiles that never cross the regression threshold, $g(T(t)) = 0$ for all $t$, and $f(T)$ is replaced by its annual mean $\bar{f}$:

$$R_0 = R_0^* \left[ 1 + \frac{\Gamma}{\bar{f}} \mathcal{I} \right].$$

### 3 Reduction to the non-structured model in the fast-progression limit

In the main text, we showed that the temperature-dependent basic reproduction number satisfies $R_0 \to R_0^*$ as $f(T) \to \infty$. Here we show that, in the same limit, the full dynamics reduce to those of the original model without explicit disease progression. The key reason is that when within-host progression is very fast, infected hosts spend negligible time in the non-chronic compartments, so transmission is effectively mediated only by the chronic-infected class.

We consider the case of constant temperature, for which $\chi(t) = 0$ and $\gamma(t) = \gamma = f(T)/\Delta\mathcal{T}$ is constant. For large $n$, progression through the infected compartments behaves as a transport process in stage space. In particular, for $k = 2, \ldots, n-1$, Eq. (3) in the main text becomes

$$\dot{I}_{H,k}(t) = \gamma \left( I_{H,k-1}(t) - I_{H,k}(t) \right). \tag{3.1}$$

Since the characteristic delay between adjacent compartments is $1/\gamma$, we expand $I_{H,k}(t + 1/\gamma)$ around $t$,

$$I_{H,k}\left(t + \frac{1}{\gamma}\right) = I_{H,k}(t) + \frac{1}{\gamma} \dot{I}_{H,k}(t) + \mathcal{O}\left(\frac{1}{\gamma^2}\right). \tag{3.2}$$

Substituting Eq. (3.1) into Eq. (3.2) yields

$$I_{H,k-1}(t) \approx I_{H,k}\left(t + \frac{1}{\gamma}\right),$$

or equivalently,

$$I_{H,k}(t) \approx I_{H,k-1}\left(t - \frac{1}{\gamma}\right). \tag{3.3}$$

Thus, consecutive infected compartments are approximately time-shifted copies of the same traveling wave.

The first infected compartment can be approximated similarly. From Eq. (2) in the main text, the exact solution is

$$I_{H,1}(t) = \beta \int_0^t S_H(s) I_v(s) e^{-\gamma(t-s)} \, ds.$$

For large $\gamma$, the exponential kernel is sharply concentrated near $s = t$. Assuming that $S_H(s)I_v(s)$ varies slowly on the timescale $1/\gamma$, we obtain

$$I_{H,1}(t) \approx \frac{\beta}{\gamma} S_H(t) I_v(t).$$

Combining Eq. (3.3) and Section 3, we arrive at

$$I_{H,k}(t) \approx \frac{\beta}{\gamma} S_H\left(t - \frac{k-1}{\gamma}\right) I_v\left(t - \frac{k-1}{\gamma}\right), \qquad k = 1, \ldots, n-1. \tag{3.4}$$

This approximation is illustrated in Fig. 1: before shifting, the compartment trajectories are displaced in time; after plotting them against $t - (k-1)/\gamma$, they collapse onto the same underlying wave.

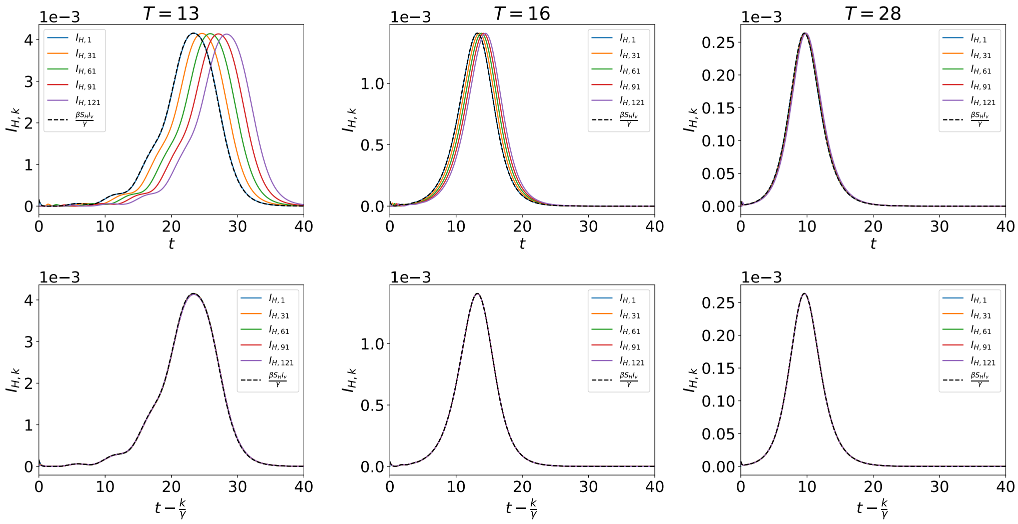


**Figure 1: Infected compartments behave approximately as a traveling wave in stage space.** First row: trajectories of selected infected compartments, $I_{H,k}$, together with the approximation $\beta S_H I_v/\gamma$. Second row: the same trajectories plotted against the shifted time $t - (k-1)/\gamma$. The collapse of the curves after this shift shows that consecutive infected compartments are approximately delayed copies of one another, consistent with the transport approximation used in the reduction.

Using $\Delta\mathcal{T} = \mathcal{T}^*/n$, we can write

$$\frac{1}{\gamma} = \frac{\mathcal{T}^*}{n\, f(T)}.$$

Therefore, the characteristic delay required for an infected host to reach the chronic compartment is

$$\tau_c = \frac{n-1}{\gamma n}\mathcal{T}^* = \frac{n-1}{n}\frac{\mathcal{T}^*}{f(T)} \approx \frac{\mathcal{T}^*}{f(T)}.$$

Substituting Eq. (3.4) into the model equations gives the approximate delayed system

$$\begin{aligned}
\dot{S}_H(t) &= -\beta S_H(t) I_v(t), \\
\dot{I}_{H,n}(t) &= \beta S_H(t-\tau_c) I_v(t-\tau_c) - \Gamma I_{H,n}(t), \\
\dot{R}_H(t) &= \Gamma I_{H,n}(t), \\
\dot{S}_v(t) &= \delta - S_v(t)\left[\sum_{i=1}^{n-1} \alpha_i \frac{\beta}{\gamma} S_H\left(t - \frac{i}{\gamma}\right) I_v\left(t - \frac{i}{\gamma}\right) + \alpha I_{H,n}(t)\right] - \mu S_v(t), \\
\dot{I}_v(t) &= S_v(t)\left[\sum_{i=1}^{n-1} \alpha_i \frac{\beta}{\gamma} S_H\left(t - \frac{i}{\gamma}\right) I_v\left(t - \frac{i}{\gamma}\right) + \alpha I_{H,n}(t)\right] - \mu I_v(t).
\end{aligned}$$

These equations clarify the role of temperature. The relevant timescale is $\tau_c \sim \mathcal{T}^*/f(T)$, which measures how long infected hosts take to reach the chronic compartment. When $\tau_c$ is large, non-chronic infected hosts contribute substantially to transmission. When $\tau_c$ is small, hosts become effectively chronic almost immediately after infection, and the delayed contributions from non-chronic stages lose weight. In the limit $f(T) \to \infty$, $\tau_c \to 0$, and the dynamics reduce to those of the original temperature-independent model. This is exactly what is observed in Fig. 2, where the trajectories of the full model converge to those of the original model as $T$ approaches $T_{\text{opt}}$.

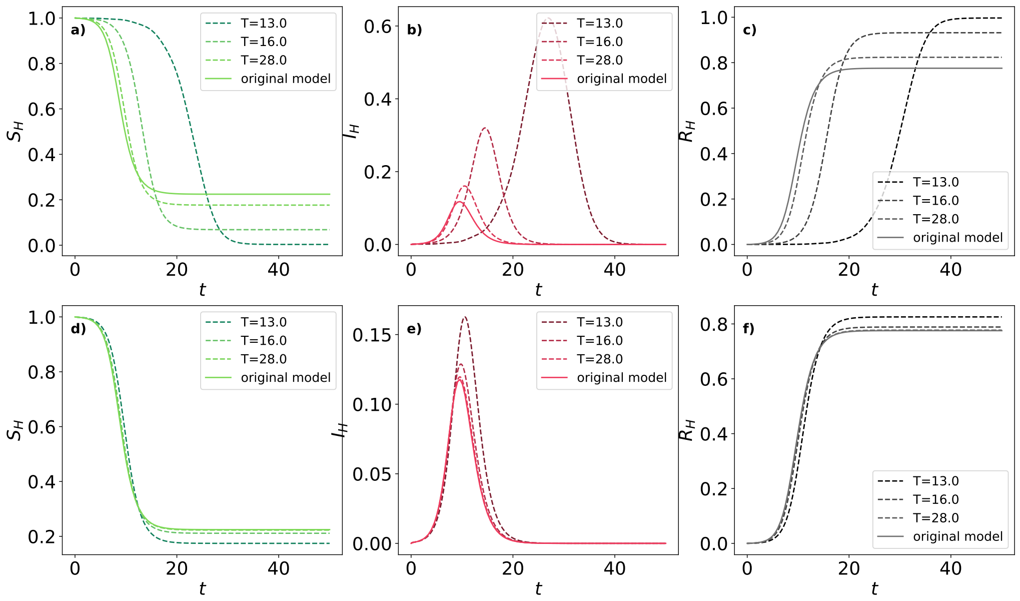


**Figure 2: The full model converges to the original model in the fast-progression limit.** Host trajectories under the stage-structured model (dashed lines) and the original model without explicit disease progression (solid lines) for three constant temperatures. Panels (a–c) show a parameter set with a larger epidemic size, and panels (d–f) a parameter set with a smaller epidemic size. In both cases, the agreement improves as temperature approaches the within-host optimum, illustrating that when progression becomes sufficiently fast, the contribution of non-chronic infected stages becomes negligible.

### 4 Temperature series

Many of the results in the main text were obtained using sinusoidal temperature forcing. This choice is motivated by the fact that empirical temperature series are dominated by a strong seasonal component, with shorter-term fluctuations superimposed on it. Here we compare the empirical series used in Fig. 5 of the main text with their fitted sinusoidal profiles, and show how the differences between both propagate into the early epidemic dynamics.

As shown in Fig. 3 **(a)**–**(d)**, the fitted sinusoidal curves capture the dominant annual cycle of the empirical temperature series, although they do not reproduce the short-term fluctuations or the exact extrema. The first issue does not have a particularly strong impact on the dynamics. Indeed, the largest departures from the sinusoidal fit arise from hourly temperature variations, which are averaged over each day to compute the corresponding daily contribution to $\mathcal{T}$, the quantity that actually governs the dynamics. Consequently, as can be seen in plots **(e)**–**(h)**, the agreement between the empirical and sinusoidal profiles is considerably better at this level.

The second issue, however, has a strong impact on the dynamics, especially at low temperatures. Most sinusoidal fits have an associated $T_{\text{min}}$ above $T_r$, implying the absence of disease regression. In contrast, as shown in Fig. 3, regression does occur for the empirical temperature series. This leads to a completely different dynamical picture, as illustrated in Fig. 5 in the main text.

To illustrate how these differences affect the dynamics, Fig. 4 compares the distribution of infected hosts across infection stages for the empirical and fitted sinusoidal temperature series at three representative times during the early phase of the epidemic.

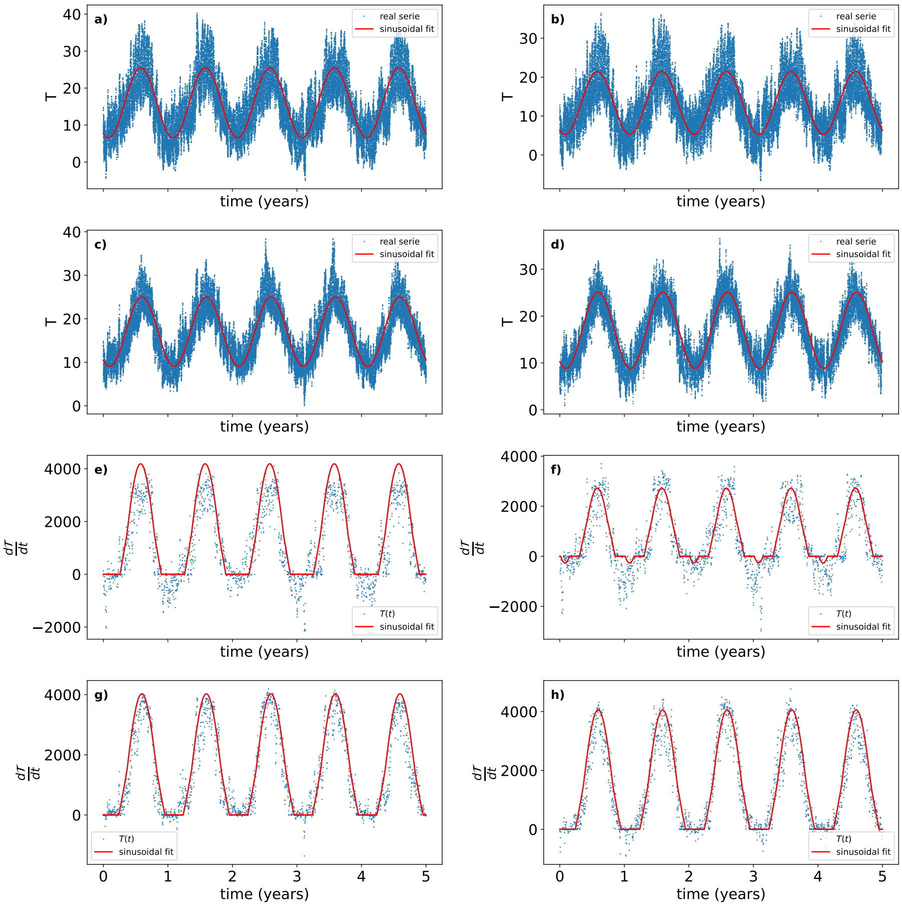


**Figure 3: Empirical temperature series are well described by a dominant seasonal component with superimposed fluctuations.** In **(a)**–**(d)**, dots represent the empirical temperature series corresponding to panels **(a)**–**(d)** of Fig (5) in the main text, while the red solid curves show the corresponding fitted sinusoidal profiles. In **(e)**–**(h)**, we show the function $\frac{d\mathcal{T}}{dt}$ corresponding to the empirical and sinusoidal temperature profiles displayed in **(a)**–**(d)**. Panel **(e)** corresponds to the temperature profiles in **(a)**, **(f)** to **(b)**, **(g)** to **(c)**, and **(h)** to **(d)**.

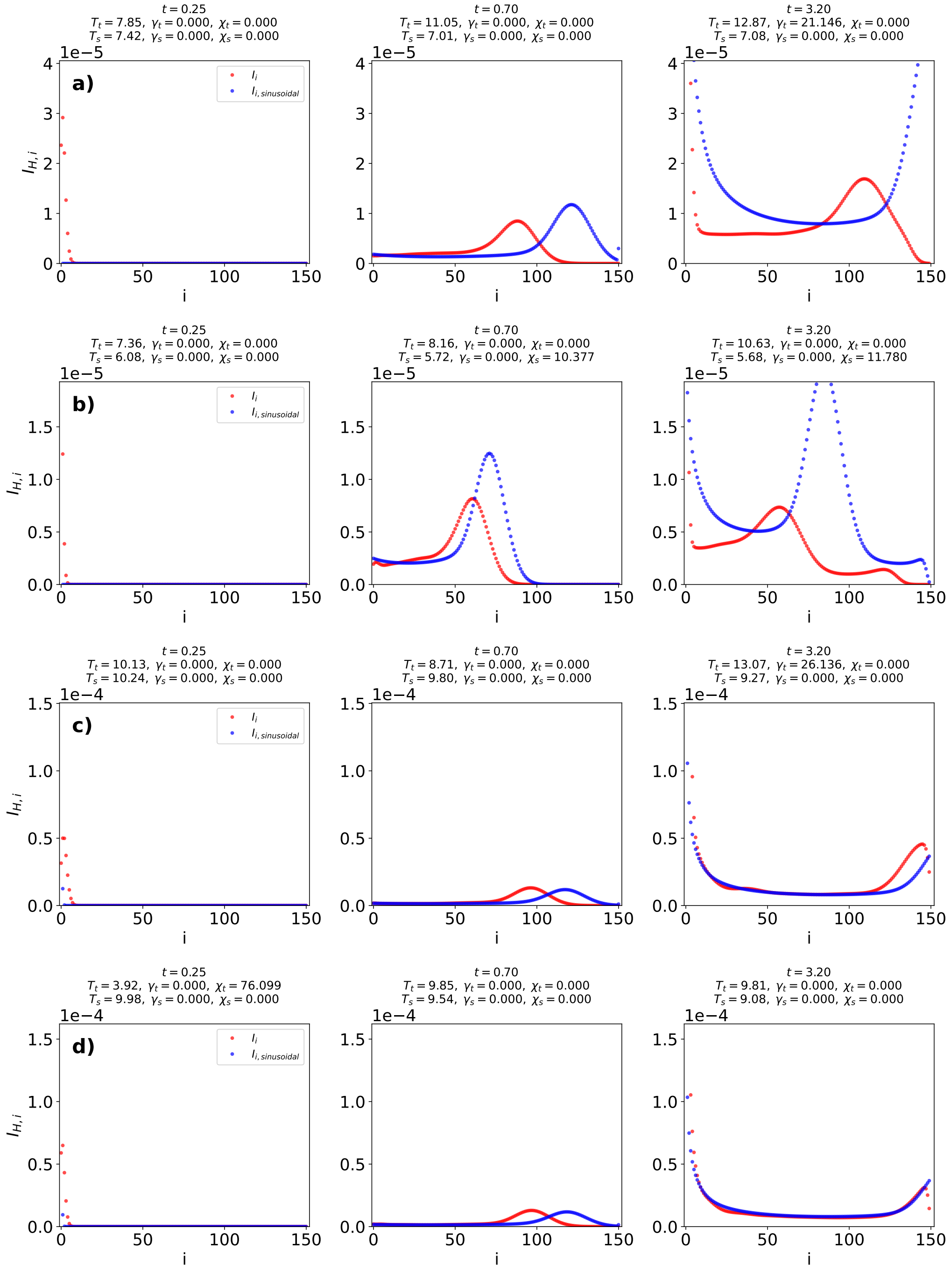


**Figure 4: Differences between empirical and sinusoidal temperature series translate into differences in early epidemic progression.** Values of $I_{H,i}$ at three representative times for the empirical temperature series and their corresponding fitted sinusoidal profiles are shown in Fig. 3.

In general, the empirical temperature series trigger earlier movement of infected hosts through the infection stages than the fitted sinusoidal curves, because short-term fluctuations can temporarily raise temperatures above $T_{\text{base}}$ even when the seasonal baseline remains below that threshold. This effect is visible, for example, in Fig. 3 **(a)** and **(e)**, where positive temperature excursions early in the year initiate disease progression before the sinusoidal fit would predict it.

At later times, however, the opposite pattern often emerges. The wave of infected hosts associated with the empirical temperature series tends to become delayed relative to the sinusoidal case and to involve fewer infected individuals. This occurs because fluctuations around the seasonal baseline favor regression over progression as temperatures approach the regression threshold $T_r$. Under those conditions, some temperature excursions fall below $T_r$, triggering recovery events and reducing the number of infected hosts that continue to advance through the infection stages.

These effects accumulate over time. In cases **(a)** and **(b)**, after three years, the distribution of infected hosts retains a similar shape for both the sinusoidal and empirical temperature series. In contrast, in cases **(c)** and **(d)**, the corresponding distributions are completely different. This is consistent with the main-text analysis of climatic variability: the impact of fluctuations is weakest when the seasonal profile remains far from the regression threshold, and strongest when fluctuations repeatedly push the system across it. Thus, the sinusoidal approximation captures the dominant mechanism underlying the epidemic dynamics, while the empirical series shows how short-term variability modulates that mechanism in practice.

### 5 Final size heatmaps

In the main text, we summarized the relationship between the temperature-dependent reproduction number and the final epidemic size using scatter plots. Here we present the same results as heatmaps over the $(T_{\min}, T_{\max})$ climate space, which makes it possible to identify the regions where invasion occurs and to compare them directly with the corresponding values of $R_0$.

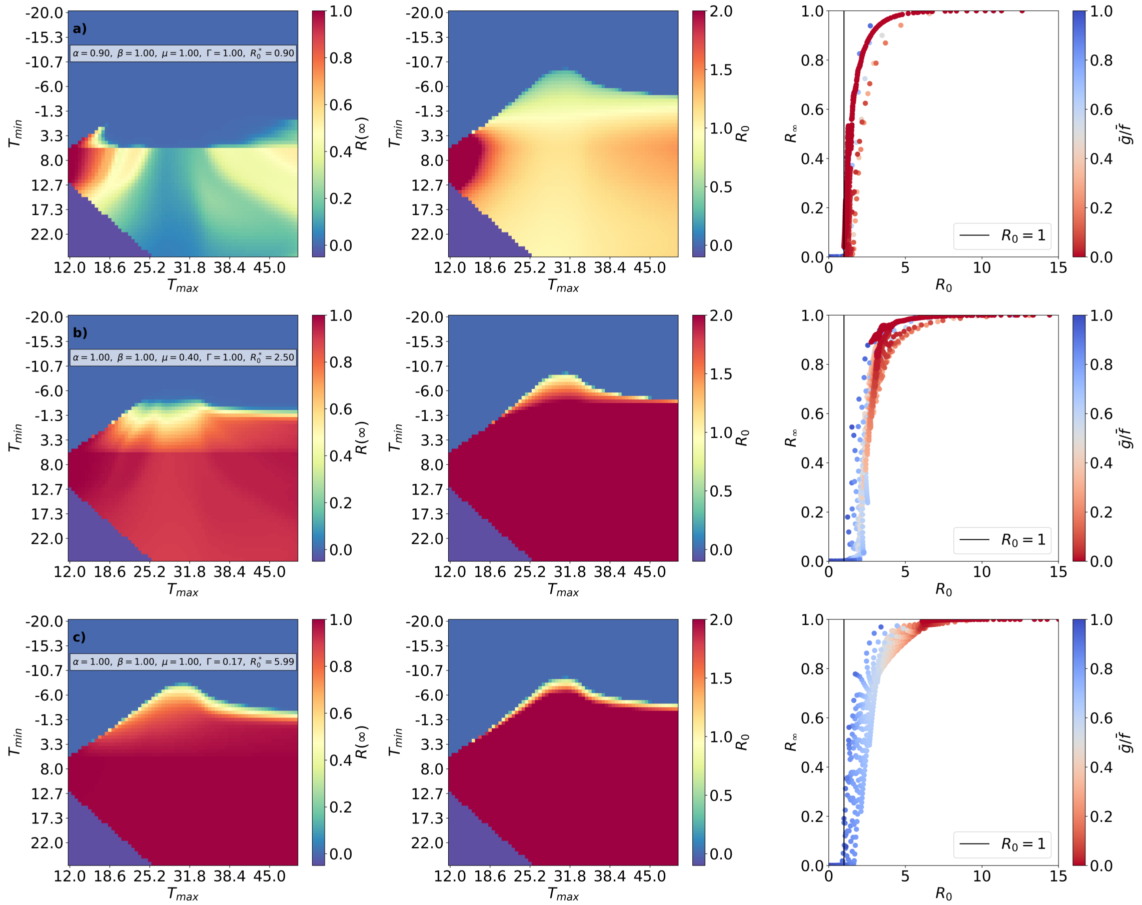


**Figure 5: Climate space associated with epidemic invasion and final epidemic size.** First column: final epidemic size, $R_H(\infty)$, across the $(T_{\min}, T_{\max})$ plane for different values of $R_0^*$. Second column: corresponding values of the temperature-dependent reproduction number. Third column: $R_H(\infty)$ plotted against the corresponding reproduction number for the same simulations.

Fig. 5 shows how the region of climate space that permits epidemic invasion changes with $R_0^*$.

As $R_0^*$ increases, the region where $R_H(\infty) > 0$ expands toward lower values of $T_{\min}$ and toward values of $T_{\max}$ closer to the thermal optimum for within-host pathogen growth. This reflects that larger values of the temperature-independent reproduction number can compensate for thermal conditions that would otherwise suppress invasion, either because progression is too fast or because regression becomes important.

The heatmaps also make clear that climatic effects can generate epidemics even when the temperature-independent system is subcritical. In the rows with $R_0^* < 1$, there are still regions of the $(T_{\min}, T_{\max})$ plane where $R_H(\infty) > 0$ and the corresponding temperature-dependent reproduction number exceeds one. These climates are characterized by a sufficiently slow progression to extend the infectious residence time while still allowing infected hosts to progress through the stages of infection.

Another prominent feature is the sharp boundary delimiting the region where epidemic invasion is possible. This boundary is closely associated with the condition $\bar{g}/\bar{f} < 1$, that is, with climates in which average progression exceeds average regression. As discussed in the main text, the approximation $R_{0,\mathrm{av}}$ is least accurate near this boundary, where small changes in temperature can shift the balance between progression and recovery and qualitatively alter the epidemic outcome. Accordingly, the misclassifications in the third column cluster near the transition between invasion and epidemic failure.

Overall, these heatmaps complement the scatter plots in the main text by explicitly showing how the same final epidemic size can arise from distinct thermal regimes and by identifying the climatic regions where the temperature-dependent contribution to invasion is strongest.

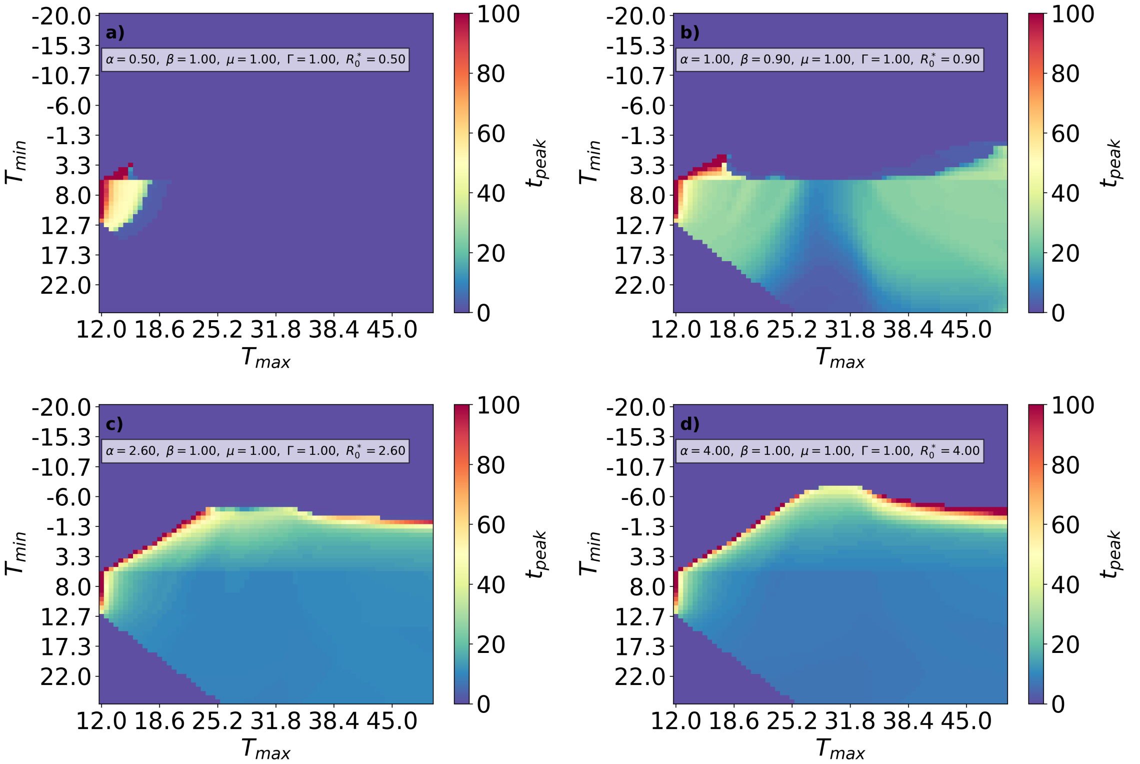


**Figure 6: Climate strongly influences the timing of epidemic development.** Values of $t_{\mathrm{peak}}$ across the $(T_{\min}, T_{\max})$ plane for different values of $R_0^*$. Blank regions correspond to parameter combinations for which no epidemic develops.

### 6 Epidemic peak time

The main text focuses on epidemic invasion and the final epidemic size. A complementary question is how climate influences the timing of epidemic development once invasion occurs. To address this, we analyze the epidemic peak time, $t_{\text{peak}}$, defined as the time at which the total density of infected hosts reaches its maximum.

Fig. 6 shows that climates allowing invasion when $R_0^* < 1$ are associated with particularly long epidemic timescales. In these cases, epidemics occur only in a restricted region of climate space where disease progression is sufficiently slow to increase transmission through prolonged residence times in infectious stages. As a consequence, the resulting epidemics develop very slowly and may require many years to reach their peak.

When $R_0^* > 1$, the dependence of $t_{\text{peak}}$ on climate is weaker over most of the $(T_{\min}, T_{\max})$ plane. The main exception occurs near the boundary separating climates that allow invasion from those that do not. As this boundary is approached, epidemics become progressively milder and slower, and $t_{\text{peak}}$ increases sharply. This is consistent with the general expectation that near the invasion threshold, epidemic growth becomes arbitrarily slow.

To relate these patterns to the thermal metrics introduced in the main text, we plot $t_{\text{peak}}$ against the inverse net progression rates.

In Fig. 7 **a)**, we observe that the peak time increases with $R_0$. For values of $R_0^*$ less than 1, there is a sharp increase in $t_{\text{peak}}$ as $R_0$ approaches 1, after which an approximately linear relationship between $R_0$ and $t_{\text{peak}}$ emerges. For $R_0^* > 1$, this sharp increase is not observed; instead, we find approximately linear growth throughout. In both cases, the slope changes at the boundary between moderate and large $R_0$ values, with a slightly higher slope in the moderate regime. Thus, in all cases, for values of $R_0$ slightly higher than 1, the relationship between $t_{\text{peak}}$ and $R_0$ can be approximated as linear.

We can understand the fact that $t_{\text{peak}}$ increases with $R_0$ by recalling that, in this regime, $R_0 \propto 1/\bar{f}$ (see the main text), and that $\bar{f}$ measures how fast the disease progresses within the host. A slower disease progression is therefore reflected in a slower epidemic progression, and vice versa.

In Fig. 7 **b)**, we generally observe higher values of $t_{\text{peak}}$ and lower values of $R_0$. This is consistent with the discussion in the main text: the effect of regression on the disease acts, on the one hand, by removing infected hosts through recovery, thereby lowering $R_0$, and, on the other hand, by delaying progression between compartments. The peak time is especially high for values of $R_0$ near 1; in this region, as discussed, epidemics persist for long times and are mild. However, in general, we do not find a clear relationship between $t_{\text{peak}}$ and $R_0$, as observed in **a)**. The case in which regression of the disease occurs is more complex, and, as discussed in the main text, it is precisely in these cases that the expression for $R_0$ is less accurate.

Fig. 7 confirms that slower average disease progression leads to delayed epidemic peaks. In the absence of regression (Fig. 7a), $t_{\text{peak}}$ increases approximately in proportion to $1/\bar{f}$ over a broad range of parameters. In this regime, the same temperatures that increase the contribution of non-chronic infectious stages to $R_0$ also delay the accumulation of infected hosts, so that larger epidemics tend to occur later.

When regression is present (Fig. 7b), the relationship becomes more dispersed, but the same general tendency remains: $t_{\text{peak}}$ increases as the net progression rate $\bar{f} - \bar{g}$ decreases. In this regime, however, slower epidemics are not necessarily larger because increased regression also reduces the number of infected hosts that progress through the stages of infection. Thus, while $t_{\text{peak}}$ remains controlled by the effective rate of disease development, its relationship with epidemic size depends on whether regression is absent or contributes substantially to the dynamics.

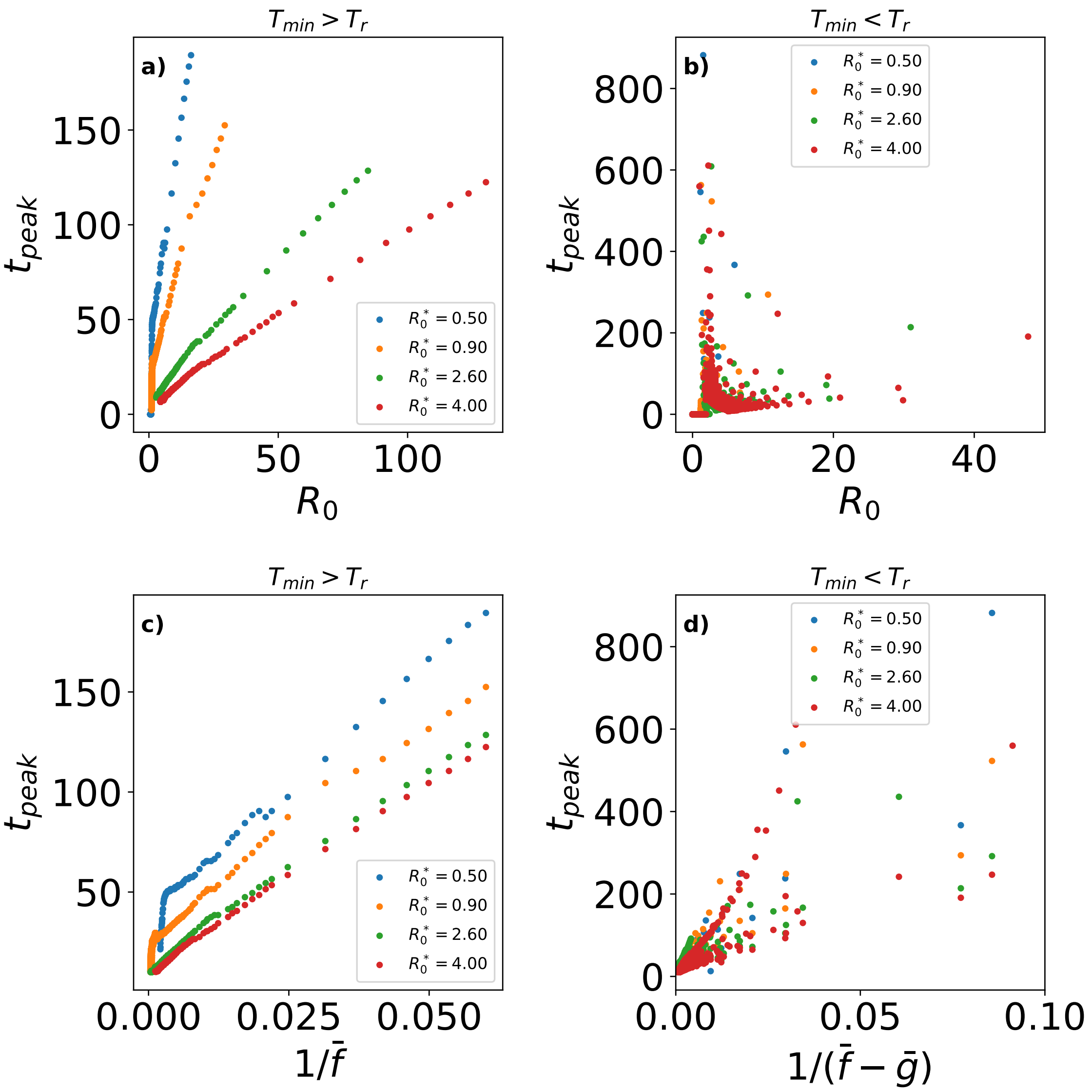


**Figure 7: Peak time increases as the effective rate of disease progression decreases.** (a) $t_{\text{peak}}$ as a function of $R_0$ for climates in which temperatures remain above the regression threshold, so that $g(T(t)) = 0$ for all $t$. (b) $t_{\text{peak}}$ as a function of $R_0$ for climates in which regression occurs during part of the year. (c) $t_{\text{peak}}$ as a function of $1/\bar{f}$ for climates in which temperatures remain above the regression threshold, so that $g(T(t)) = 0$ for all $t$. (d) $t_{\text{peak}}$ as a function of $1/(\bar{f} - \bar{g})$ for climates in which regression occurs during part of the year. Colors indicate different values of $R_0^*$.

### 7 Final size with noise heatmaps

To complement the analysis in the main text, we examined how short-term temperature variability modifies the final epidemic size across the $(T_{\text{min}}, T_{\text{max}})$ climate space for different values of $R_0^*$ and noise intensity $\sigma^2$.

Fig. 8 shows that the effect of noise is strongly heterogeneous across climate space. In most cases, variability either leaves the final epidemic size nearly unchanged or reduces it, in agreement with the main-text result that short-term fluctuations tend to suppress epidemic invasion by favoring regression. The strongest reductions occur near the climatic transition between epidemic spread and failure, where small temperature perturbations can qualitatively alter the balance between progression and recovery.

To summarize these patterns across parameter combinations, Table 1 reports the fraction of climates in which noise decreases, increases, or leaves nearly unchanged the final epidemic size, together with the mean and maximum magnitude of those changes.

**Table 1: Summary statistics of the effect of temperature variability on final epidemic size.** For each combination of $R_0^*$ and $\sigma^2$, $f_-$, $f_+$, and $f_=$ denote the fractions of climates for which noise decreases, increases, or leaves approximately unchanged the final epidemic size, respectively. The remaining columns report the mean and maximum magnitude of the negative and positive changes in $R_H(\infty)$.

| | $f_-$ | $f_+$ | $f_=$ | $\overline{\Delta R_H(\infty)_-}$ | $\max(\Delta R_H(\infty)_-)$ | $\overline{\Delta R_H(\infty)_+}$ | $\max(\Delta R_H(\infty)_+)$ |
|---|---|---|---|---|---|---|---|
| $R_0^* = 2.5,\ \sigma^2 = 2.25$ | 0.002 | 0.222 | 0.776 | 0.819 | 1.000 | 0.031 | 1.000 |
| $R_0^* = 2.5,\ \sigma^2 = 9.0$ | 0.015 | 0.310 | 0.675 | 0.116 | 0.997 | 0.068 | 1.000 |
| $R_0^* = 2.5,\ \sigma^2 = 20.25$ | 0.096 | 0.346 | 0.558 | 0.026 | 0.977 | 0.104 | 1.000 |
| $R_0^* = 0.9,\ \sigma^2 = 2.25$ | 0.038 | 0.187 | 0.776 | 0.056 | 1.000 | 0.069 | 1.000 |
| $R_0^* = 0.9,\ \sigma^2 = 9.0$ | 0.137 | 0.265 | 0.598 | 0.027 | 0.928 | 0.120 | 1.000 |
| $R_0^* = 0.9,\ \sigma^2 = 20.25$ | 0.166 | 0.293 | 0.541 | 0.039 | 0.729 | 0.157 | 1.000 |
| $R_0^* = 5.0,\ \sigma^2 = 2.25$ | 0.002 | 0.186 | 0.812 | 0.852 | 1.000 | 0.024 | 1.000 |
| $R_0^* = 5.0,\ \sigma^2 = 9.0$ | 0.002 | 0.273 | 0.725 | 0.728 | 1.000 | 0.054 | 1.000 |
| $R_0^* = 5.0,\ \sigma^2 = 20.25$ | 0.008 | 0.318 | 0.674 | 0.220 | 0.996 | 0.088 | 1.000 |

Two features stand out in Table 1. First, the maximum reduction in $R_H(\infty)$ is close to one in nearly all cases. This indicates that there are climates for which a severe epidemic under smooth seasonal forcing can be almost completely suppressed once short-term variability is introduced. As shown in the main text, these climates lie predominantly near the boundary where progression and regression nearly balance, so that small perturbations in the temperature profile can switch the system from epidemic spread to epidemic failure.

Second, increases in $R_H(\infty)$ are generally much smaller than decreases, except when $R_0^*$ is already close to the invasion threshold. This asymmetry reflects the structure of the temperature response functions $f(T)$ and $g(T)$. Noise can shift temperatures away from the within-host optimum and thereby prolong the infectious residence time, but in much of climate space it also increases regression, which counteracts or dominates that effect. Consequently, variability more often weakens epidemics than strengthens them.

A related question is how noise modifies the average progression and regression rates themselves. Since the climatic contribution to invasion depends on $\bar{f}$ and $\bar{g}$, their response to stochastic fluctuations provides a direct explanation for the patterns observed in Fig. 8. Using Jensen's inequality, one can show that if $\varphi$ is convex, then

$$\varphi(\mathbb{E}[X]) \leq \mathbb{E}[\varphi(X)],$$

whereas if $\varphi$ is concave the inequality is reversed:

$$\varphi(\mathbb{E}[X]) \geq \mathbb{E}[\varphi(X)].$$

For $X = T + \delta$, with $\mathbb{E}[\delta] = 0$, this implies

$$\mathbb{E}[X] = T.$$

Therefore, if $\varphi$ is convex,

$$\varphi(T) \leq \mathbb{E}[\varphi(T + \delta)],$$

whereas if $\varphi$ is concave,

$$\varphi(T) \geq \mathbb{E}[\varphi(T + \delta)].$$

In our case, $g(T)$ is convex over its non-zero domain, so adding noise can only increase, or leave unchanged, its mean value $\bar{g}$. By contrast, $f(T)$ is piecewise linear and changes curvature across its domain, so the effect of noise on $\bar{f}$ depends on the part of the thermal niche explored by the seasonal profile. This local application of Jensen's inequality explains why variability systematically enhances regression, while its effect on progression may be positive or negative depending on the underlying climate.

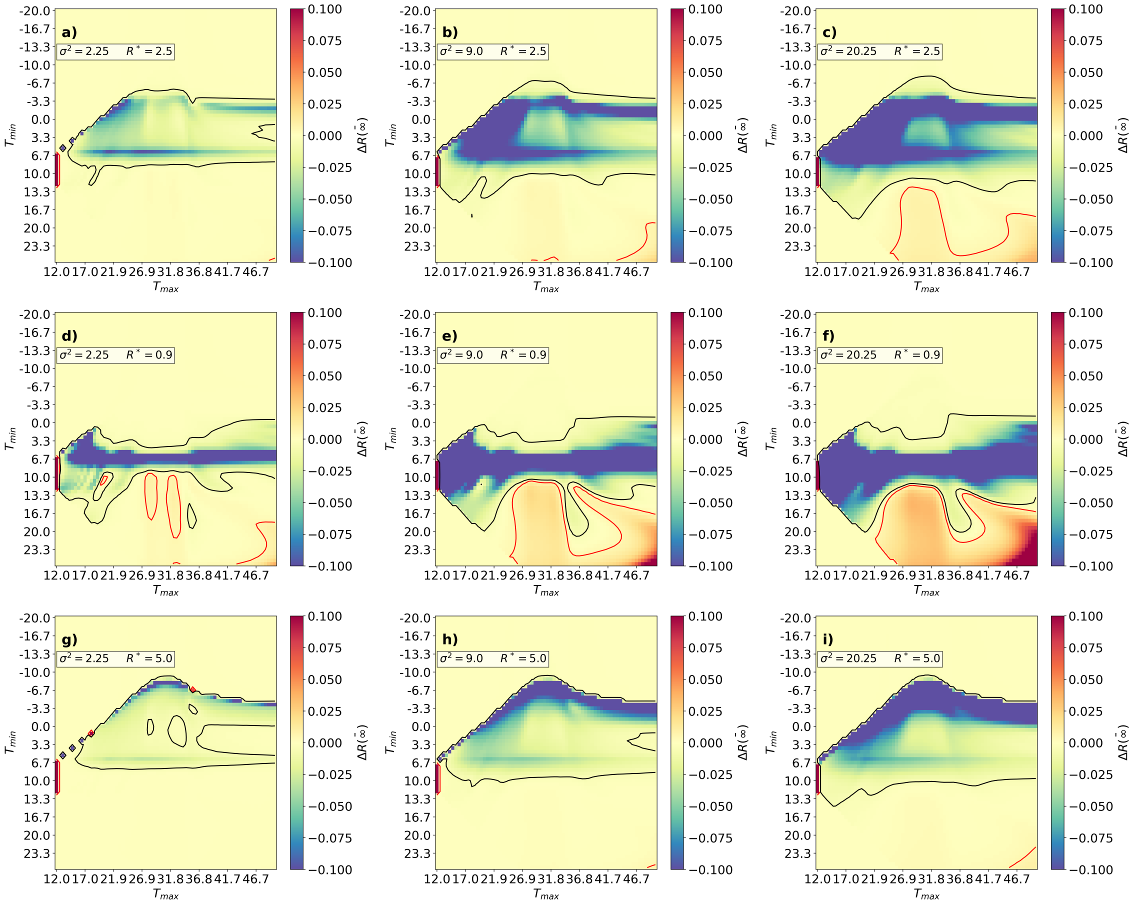


**Figure 8: Short-term climatic variability most strongly alters epidemic size near thermal thresholds.** Change in final epidemic size, $\Delta R_H(\infty)$, across the $(T_{\min}, T_{\max})$ plane for different values of $R_0^*$ and noise intensity $\sigma^2$. For each parameter combination, $\Delta R_H(\infty)$ was computed as the difference between the mean final epidemic size across 100 noisy realizations and the corresponding value obtained for $\sigma = 0$.

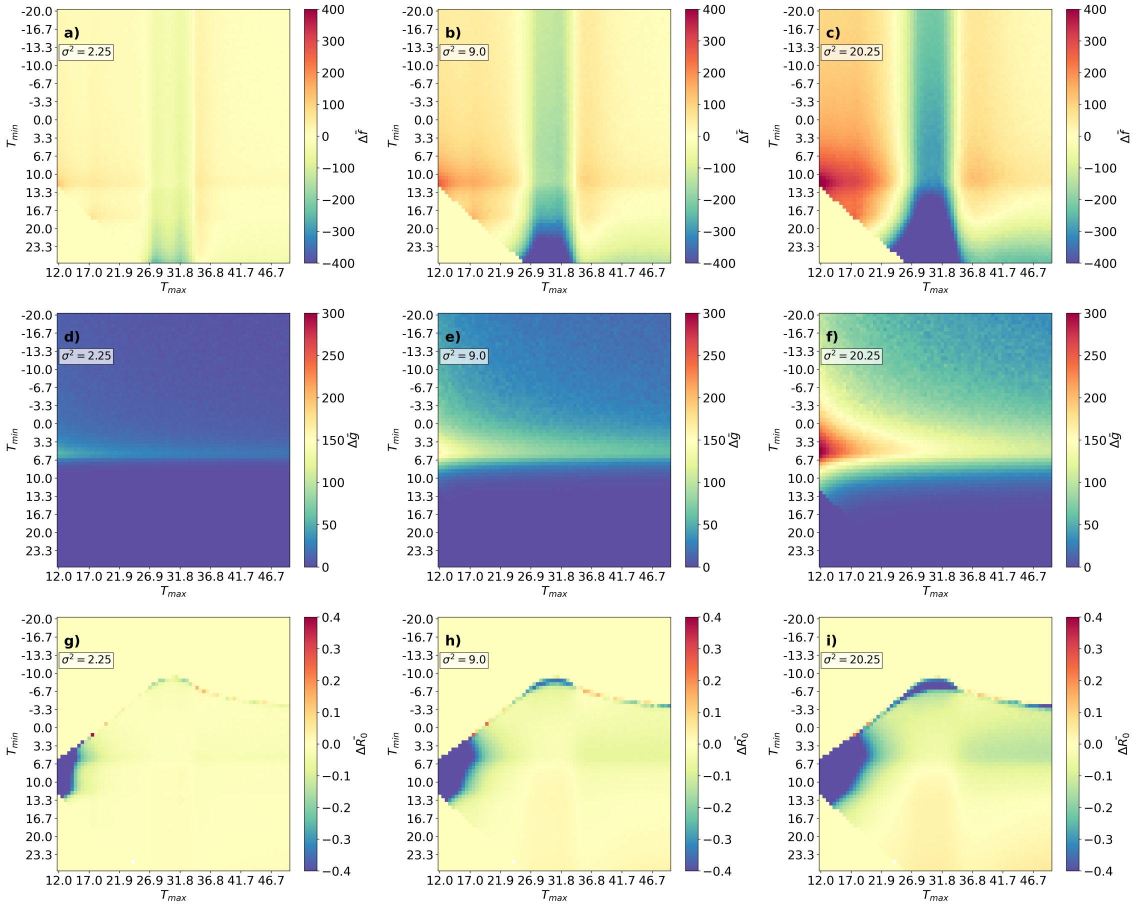


**Figure 9: Noise affects progression and regression asymmetrically.** Changes in the mean progression rate ($\Delta\bar{f}$), mean regression rate ($\Delta\bar{g}$), and normalized temperature-dependent contribution to invasion ($\Delta R_0/R_0^*$) as functions of $\sigma^2$. For each value of $\sigma^2$, the plotted quantities were averaged over 100 realizations and compared with the corresponding values for $\sigma = 0$.

This asymmetry is illustrated in Fig. 9. As noise intensity increases, $\bar{g}$ consistently increases, while $\bar{f}$ can either increase or decrease depending on the underlying temperature regime. For low levels of variability, fluctuations often move temperatures into ranges where $f(T)$ is larger, producing an increase in $\bar{f}$. For stronger variability, however, more temperatures are displaced away from the favorable part of the thermal niche, and $\bar{f}$ declines. Together, these effects explain why noise tends to reduce epidemic invasion over most of climate space, while only weakly enhancing it in a comparatively small subset of climates.

### 8 Site-specific temperature series parameters

Table 2 reports the key quantities derived from the six empirical temperature series used in Fig. 5 of the main text.

| | Sierra de Gata | Pegarinhos | Santa Maria del Camí | Bari | Castelo de Marvão | Fundão |
|---|---|---|---|---|---|---|
| $\bar{T}$ | 15.82 | 13.17 | 16.87 | 16.80 | 15.19 | 14.57 |
| $\sigma_T^2$ | 69.94 | 55.72 | 42.60 | 42.26 | 60.04 | 61.35 |
| $\bar{f}$ | 1285.88 | 947.84 | 1529.27 | 1564.98 | 1183.87 | 1123.94 |
| $\frac{\bar{g}}{\bar{f}}$ | 0.07 | 0.19 | 0.00 | 0.01 | 0.07 | 0.10 |
| $R_0$ | 6.53 | 6.42 | 6.69 | 6.62 | 6.72 | 6.60 |
| $T_{\min}$ | 6.39 | 5.12 | 8.92 | 8.70 | 6.86 | 6.02 |
| $T_{\max}$ | 25.32 | 21.31 | 24.88 | 24.97 | 23.60 | 23.20 |
| $\mu_r$ | 0.00 | 0.00 | 0.00 | 0.00 | 0.00 | 0.00 |
| $\sigma_r^2$ | 25.01 | 22.86 | 10.66 | 9.06 | 24.87 | 24.38 |
| lat | 40.10 | 41.30 | 39.70 | 41.10 | 39.40 | 40.10 |
| lon | -6.80 | -7.40 | 2.80 | 16.90 | -7.40 | -7.50 |

**Table 2:** Some quantities related to the temperature series corresponding to the selected points and relevant to the model are reported: latitude (lat), longitude (lon), mean temperature $\bar{T}$, its variance $\sigma_T^2$, the mean of $f(T)$, $\bar{f}$, the mean of $g(T)$ relative to the mean of $f(T)$, $\frac{\bar{g}}{\bar{f}}$, and $R_0$. We also report several quantities obtained from fitting the temperature series to a sinusoidal function: the minimum temperature of the fitted sinusoid, $T_{\min}$, the maximum temperature of the fitted sinusoid, $T_{\max}$, and the residuals $r(t) = T(t) - T_{\sin}(t)$, whose mean and variance are denoted by $\mu_r$ and $\sigma_r^2$, respectively.
